# Evaluating deep learning for predicting epigenomic profiles

**DOI:** 10.1101/2022.04.29.490059

**Authors:** Shushan Toneyan, Ziqi Tang, Peter K. Koo

**Author notes:** These authors contributed equally to this work (Shushan Toneyan, Ziqi Tang).

## Abstract

Deep learning has been successful at predicting epigenomic profiles from DNA sequences. Most approaches frame this task as a binary classification relying on peak callers to define functional activity. Recently, quantitative models have emerged to directly predict the experimental coverage values as a regression. As new models continue to emerge with different architectures and training configurations, a major bottleneck is forming due to the lack of ability to fairly assess the novelty of proposed models and their utility for downstream biological discovery. Here we introduce a unified evaluation framework and use it to compare various binary and quantitative models trained to predict chromatin accessibility data. We highlight various modeling choices that affect generalization performance, including a downstream application of predicting variant effects. In addition, we introduce a robustness metric that can be used to enhance model selection and improve variant effect predictions. Our empirical study largely supports that quantitative modeling of epigenomic profiles leads to better generalizability and interpretability.

---

Deep learning (DL) has achieved considerable success in predicting epigenomic profiles from DNA sequences, including transcription factor binding^1–3^, chromatin accessibility^4,5^, methylation^6^, and histone marks^7,8^. By learning a sequence-function relationship, trained DL models have been utilized on various downstream tasks, such as predicting the functional effects of single-nucleotide variants associated with human diseases^4,7,9–13^.

Over the past several years, the variety of DL models proposed to address regulatory genomic prediction tasks has increased substantially^14–22^. The wide variety of proposed models, the datasets they are trained on, how the datasets are processed, and the tricks used to train the models make it challenging to assess which innovations are driving performance gains. A direct comparison of model performance cannot always be made easily due to the variations of how the prediction tasks are framed. For instance, previous approaches typically frame the task as a *binary* classification, where binary labels represent functional activity based on a peak caller. However, in collapsing the amplitude and shape of a peak into a binary label, information about differential *cis*-regulatory mechanisms potentially encoded in these attributes is lost. Recently, *quantitative* models^23–27^ have emerged, similarly taking DNA sequences as input but now directly predicting experimental read coverage values as a regression task, thus bypassing the need for a peak caller and preserving quantitative information of epigenomic tracks. Since standard metrics differ across classification and regression tasks, it remains unclear how to directly compare models trained on binary tasks versus quantitative tasks.

To address this issue, Kelley et al^23^ propose to ‘binarize’ their quantitative predictions using a peak caller, which enables a comparison of the overlapping regions with binary labels. However, this approach narrowly focuses evaluation on regions of the genome that have been annotated as functional according to a peak caller, which is noisy and sensitive to parameter choices of the peak caller^28^. Alternatively, Avsec et al^26^ compared the performance of a binary model with an augmented version of the binary model that appends an output-head that simultaneously predicts quantitative profiles. While this measures the added benefit of quantitative modeling, this approach requires retraining multiple versions of the model, which can be sensitive to initialization, and it does not easily extend to comparisons with existing models.

Moreover, other modeling choices within a prediction task make it challenging to directly make fair comparisons. For instance, existing quantitative models predict different resolutions of the epigenetic profiles. Basenji^23^ predicts non-overlapping binned epigenomic profiles with a resolution of 128 base-pairs (bp), while BPNet^26^ predicts at base-resolution. Comparing models across different resolutions is not straightforward, because binning affects the smoothness of the coverage values which, in turn, can influence performance metrics. Moreover, existing methods employ different data augmentations and analyze different subsets of training and test data, further complicating any direct comparisons.

As the number of applications continues to grow, a bottleneck of modeling innovations is forming as we lack the ability to perform a critical assessment of newly proposed models. Here, we propose an evaluation framework for DL models trained on regulatory genomics data that enables a systematic comparison of prediction performance and model interpretability, irrespective of how the prediction task is framed. Using this framework, we perform a critical assessment of quantitative models and binary models on a chromatin accessibility prediction task to elucidate beneficial factors in model architecture, training procedure, and data augmentation strategies. Moving beyond predictive performance, we assess each model with additional criteria: 1) robustness of predictions to small perturbations to the input sequence, 2) variant effect predictions, and 3) interpretability of the learned representations. Our evaluation framework is packaged in a python-based software, called GOPHER (GenOmic Profile-model compreHensive EvaluatoR).

## Results

Many newly proposed DL models are accompanied with custom software; however, their scope is often limited to employing a specific pipeline, making it difficult to mix-and-match innovations across methods. To gain deeper insights into the factors that drive model performance, it is critical to be able to make a systematic and fair comparison across existing and newly proposed DL models. To address this gap, we developed a new, integrative software package called GOPHER that consists of high-level Tensorflow/Keras-based APIs for data processing, data augmentation strategies, and comprehensive model evaluation, including variant effect predictions and model interpretability, for binary and quantitative modeling of epigenomic profiles (Fig. 1).

**Figure 1.**
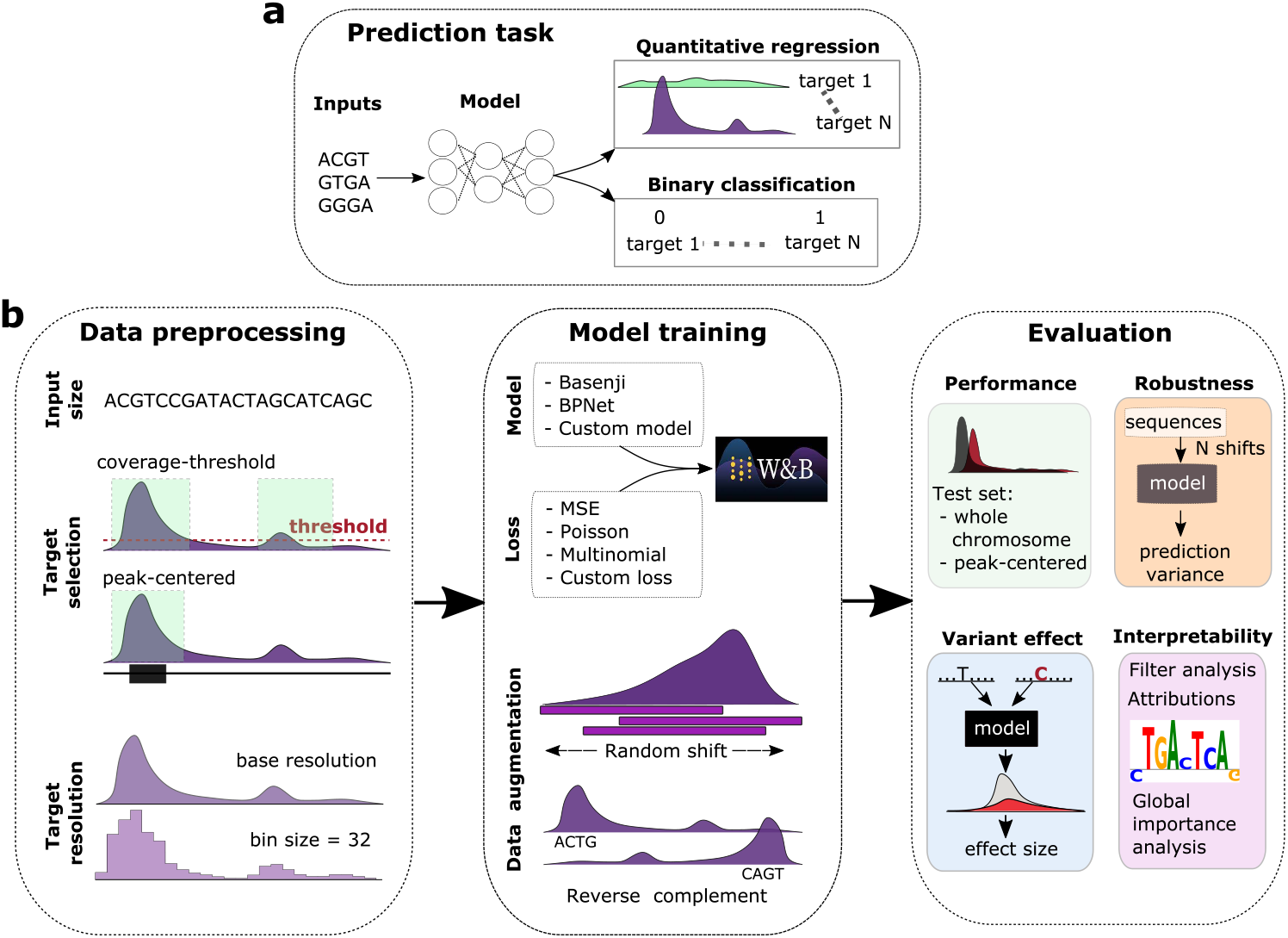
GOPHER overview. (**a**) Comparison of binary and quantitative prediction tasks for regulatory genomics. (**b**) Illustration of the 3 main components of DL analysis: data preprocessing (i.e. input size, target selection and resolution), model training (i.e. model architecture, loss and data augmentations) and evaluation (i.e. generalization performance, robustness, interpretability and variant effect predictions).

### Performance evaluation of best-in-class quantitative models

Prominent quantitative models for regulatory genomics are Basenji^23^ and BPNet^26^. Each employ different strategies for model design, data processing, loss function, evaluation metric, and data augmentations (Supplementary Table 1), which makes it difficult to identify the key factors that drive performance gains. Thus, we performed a systematic comparison of Basenji- and BPNet-inspired models on a multi-task quantitative prediction of chromatin accessibility ATAC-seq data across 15 human cell lines (see Methods). This dataset provides a sufficient challenge in deciphering the complexity of enhancer activity across cell types but maintains a dataset size that is amenable to the scale of comprehensive evaluations performed in this study.

For each base model, we used GOPHER to search for optimal hyperparameters using each model’s original *target resolution* and *training set selections* (Supplementary Fig. 1). Target resolution defines the bin size of the prediction task, which is used to create non-overlapping windows of coverage values, with the lowest resolution being a bin size of the entire input sequence (i.e. predicting a single quantitative output) while the highest resolution is a bin size of 1 (i.e. base-resolution). BPNet was trained at base resolution on *peak-centered* data (BPNet-base), which consists of a training set selection of genomic regions that contain at least 1 peak from a target cell-type. On the other hand, Basenji was trained at 128 bin-resolution (Basenji-128) with *coverage-threshold* data, which consists of training set selection based on segmenting each chromosome into non-overlapping regions and then sub-selecting the regions that have a max coverage value above a set threshold. Details of the default choices for the dataset and training parameters are detailed in Methods.

Overall, the quality of the model predictions were in line with previous studies (Fig. 2a). Using the optimized models as a baseline, we compared the impact of various factors that influence prediction performance, including loss function, target resolution, training set selection, and test set selection.

**Figure 2.**
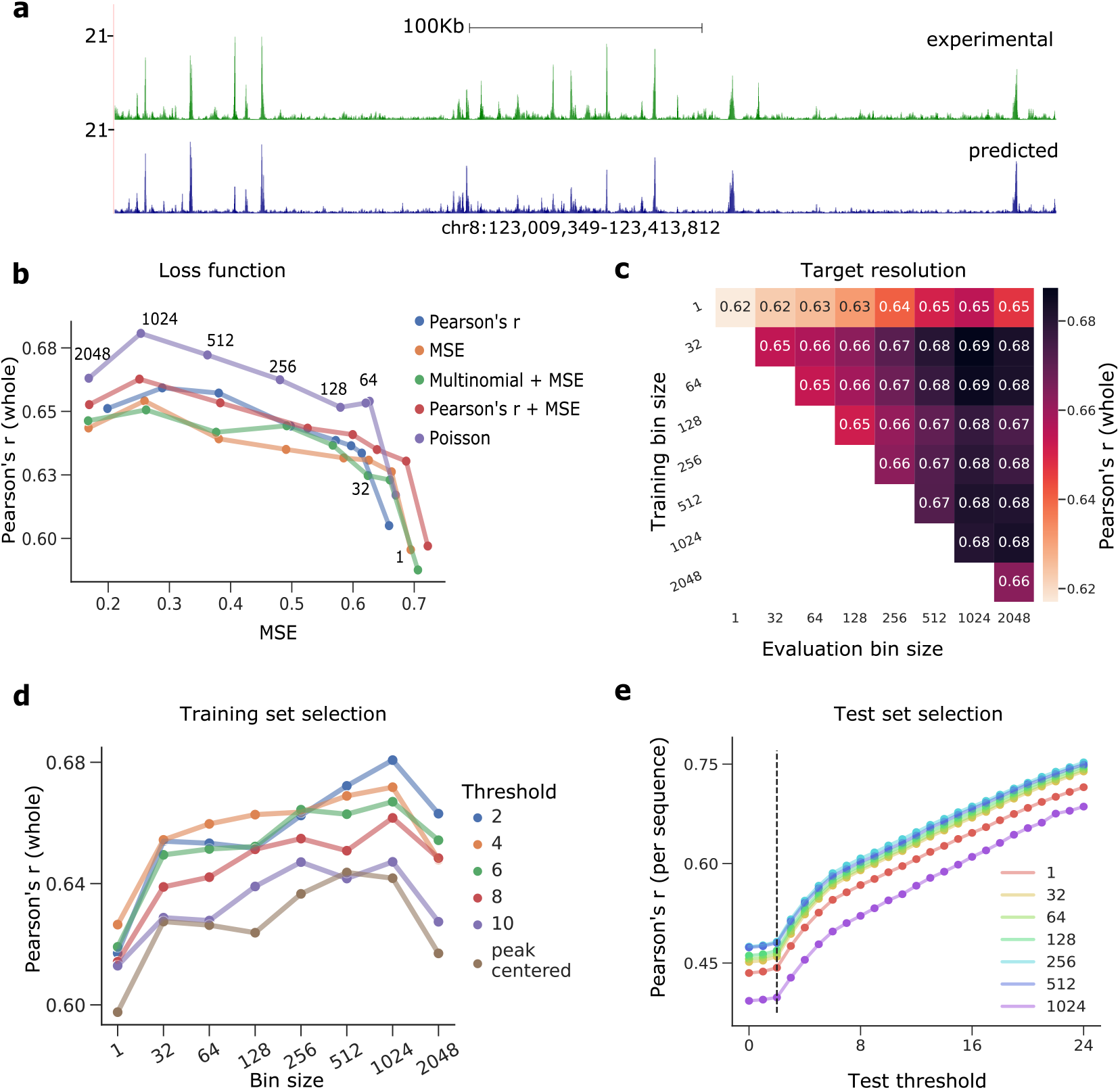
Evaluation of Basenji-based quantitative models. (**a**) Example visualization of bigWig tracks for experimental measurements (top) and model predictions (bottom) for a given target cell line (GM23338) on a held-out test chromosome for Basenji-128. (**b**) *Loss function analysis*. Scatter plot of the whole-chromosome Pearson’s r versus the MSE for different loss functions (shown in a different color) and target resolutions (annotated bin sizes). Predictions were scaled for all models using ground truth mean coverage (see Methods). Lines serve as a guide-to-the-eye. (**c**) *Target resolution analysis*. Heatmap of the whole-chromosome Pearson’s r for models trained on a given bin size (*y*-axis) with predictions that were systematically down-sampled to a lower resolution for evaluation (*x*-axis). (**d**) *Training set selection analysis*. Scatter plot of whole-chromosome Pearson’s r versus different target resolutions (i.e. bin size) for Baenji-based models trained on datasets with a different coverage threshold applied to the training set (shown in a different color). Peak-centered represents when the data is trained only on genomic regions identified as a peak. (**e**) *Test set selection analysis*. Scatter plot of the thresholded Pearson’s r, which is average of per sequence correlation in the thresholded test set, versus different coverage thresholds applied to the test set for various Basenji-based models (of different resolutions) trained on default data, i.e. coverage-threshold data with a threshold of 2 (indicated with the black dotted vertical line). (**b**-**e**) Pearson’s r represents the average across cell lines.

#### Loss function

The choice of loss function for quantitative models is not as straightforward as binary models, which is typically a binary cross-entropy. Loss functions can penalize the shapes (e.g. Pearson’s r) or the magnitudes (e.g. mean-squared-error (MSE)). BPNet employs a combination of MSE for the magnitude and multinomial loss for the shape. On the other hand, Basenji employs a single loss, Poisson negative log-likelihood (NLL). To explore the effect of loss function on quantitative modeling, we systematically evaluated Basenji-based models and BPNet-based models (using the optimized parameter setting from hyperparameter search) across 5 different loss functions at 8 different target resolutions (Fig. 2b). Evidently, Poisson NLL outperformed the other losses at all tested bin-resolutions, i.e. lower MSE and higher Pearson’s r on the held out test set. On the other hand, Pearson’s r and the combination of Pearson’s r and MSE loss yielded the second best overall performance. Interestingly, higher bin sizes tend to yield better performance up to a bin size of about 1 kb for Basenji-based models, which is roughly the width of an ATAC-seq peak. Surprisingly, BPNet-based models yielded a different trend, where base-resolution models performed the best, albeit Poisson NLL remained the best loss (Extended Data Fig. 1a). This is expected (to a degree) as each of these models were optimized for different resolutions. This suggests that model design can be optimized for a given resolution but may not necessarily generalize across resolutions.

#### Target resolution

Quantitative models that employ different target resolutions cannot be directly compared, because the bin sizes serve to down-sample the number of data points, which can affect performance metrics based on correlation and binning effectively smoothens high-frequency noise. To explore whether the observed relationship between a higher bin size (i.e. lower resolution) and higher prediction performance for Basenji-based models is due to more accurate predictions or because of the statistical artefacts from binning, we developed an evaluation scheme that enables a direct comparison across target resolutions. Specifically, we binned the predictions of the higher resolution models to match the lower resolution predictions. This effectively provides an avenue to fairly compare the performance across target resolutions. As expected, models trained at a given target resolution yield a higher Pearson’s r with increased smoothing, despite that the biology underlying the predictions remains unchanged (Fig. 2c). A similar observation was made for BPNet-based models (Extended Data Figure 1b). To further demonstrate the sensitivity of Pearson’s r on smoothness properties, we systematically smoothed predictions while maintaining the predicted resolution by applying a box-car filter with window sizes that matched a lower-resolution bin and observed a similar trend (Extended Data Figure 2).

#### Training set selection

One major component of generalization performance depends on the composition of training data. To explore the impact of training set selection, we systematically trained Basenji-based models at different resolutions on new datasets with increasing stringency of a coverage threshold, which serves to modulate the balance between the original BPNet’s peak-centered training approach and the original Basenji’s whole-chromosome training approach. For comparison, we also trained each model on peak-centered data. By evaluating each model on the whole-chromosome test set for consistency, we found that the training set with the lowest threshold yielded the best performance, while peak-centered models performed the worst (Fig. 2d). Limiting the model to only higher functional activity reduces the data set size, and hence the number of examples the model has to learn from, which may explain some of the drop in performance. On the other hand, providing too many inactive regions may imbalance the model’s focus on features within inactive sites, though we did not observe this undesirable behavior.

#### Test set selection

Choice of test set can influence the measure of generalization performance. The common approach is to process the training set and test set in the same way and split them via random splits or held-out chromosome(s). Alternatively, predictions across the whole chromosome (via tiled predictions) puts a greater emphasis on generalization to regions of non-functional sites. To explore the influence of test set selection on model performance, we generated new test sets with progressively stringent coverage thresholds to modulate between the two extremes. To compare performance across test thresholds, we calculated the average Pearson’s r per sequence, instead of calculating a single Pearson’s r across tiled predictions. Interestingly, Basenji-base’s performance monotonically increases with the coverage threshold on the test set (Fig. 2e). A similar trend was observed across other resolutions. This illustrates that predictions are more accurate for higher coverage regions and thus focusing only on high-activity regions can inflate test performance.

In addition, we performed a more targeted evaluation of the performance at high-activity regions across models trained on either peak-centered or coverage-threshold datasets (Extended Data Table 1). We observed a consistent trend where models trained on peak-centered data had a slight performance advantage over models trained on coverage-threshold data in terms of scaling the heights of the reads (i.e. lower MSE). However, models trained on coverage-threshold data yielded substantially better peak detection performance across the whole chromosome (i.e. higher Pearson’s r). Thus, there is a slight trade-off in scaling the predictions when training on coverage-threshold data, but it leads to lower false positive activity predictions genome-wide.

### Robustness test to identify models with fragile predictions

Robustness to input perturbations is a widely used criterion for evaluating the trustworthiness of DL models^29,30^. Adversarial attacks using small, targeted noise to the inputs dramatically affects the prediction of non-robust models^31^. These noise-based perturbations do not naturally extend to genomic data. Alternative perturbations, such as single-nucleotide mutations, can affect function and hence are also inappropriate. We developed a robustness test to measure the sensitivity of model predictions to translational shifts of the input sequences, whose function is largely maintained by also shifting the target predictions (see Methods). Specifically, our robustness test provides a variation score for a given input sequence that is randomly translated *N* times – the predictions are aligned and only overlapping regions are considered for the variation summary statistic (Fig. 3a).

**Figure 3.**
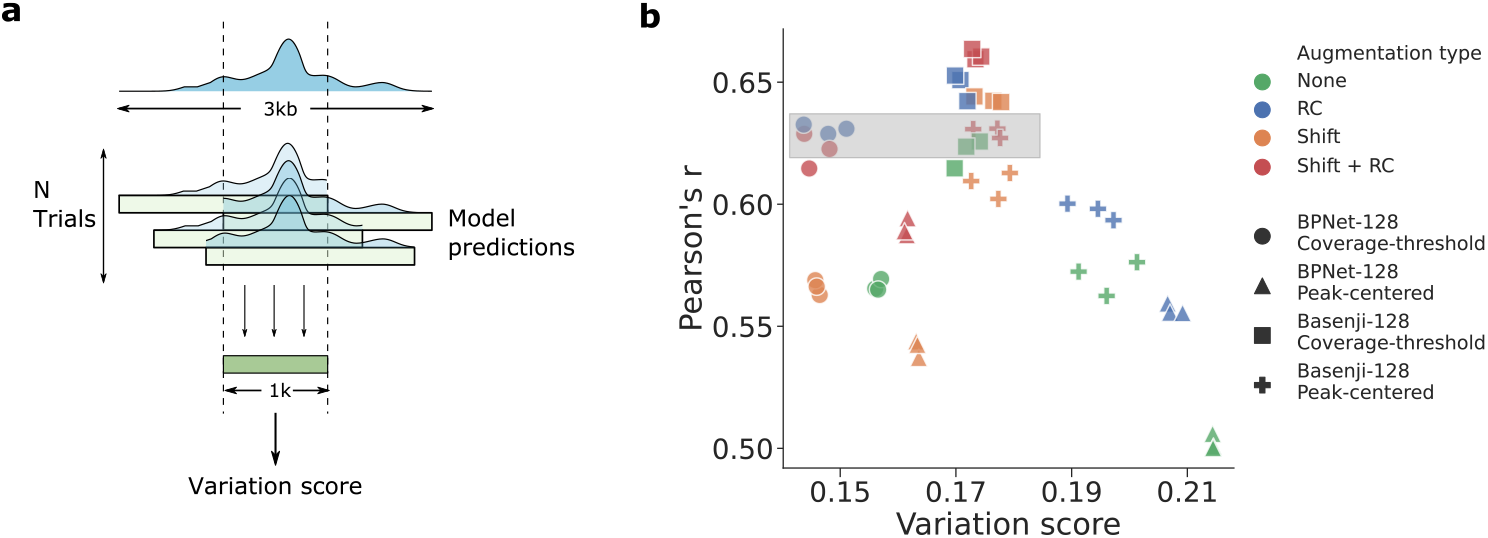
Testing model robustness against translational shifts. (**a**) Schematic overview of robustness test. For each 3 kb sequence, N random 2 kb sub-sequences were extracted, and the standard deviation across predictions within the overlapping regions is calculated. Average variation score of predictions (i.e. average per-position standard deviation of coverage values normalized by the total mean coverage value) was used as a measure of model robustness. (**b**) Scatter plot of the Pearson’s r (averaged across a per-sequence analysis) versus the robustness variation score across models with different augmentation methods (shown in a different color). Each 128 bin-resolution model (shown in a different marker) was trained 3 times with different random initializations. Pearson’s r represents the average across cell lines.

To test the robustness of models across augmentation strategies and choice of training sets, we compared how different combinations of augmentations, including random reverse-complement (RC) transformations and random shifts of the input sequence (up to 1024 bp), affect the model’s robustness properties for BPNet-128 and Basenji-128 trained on either peak-centered or coverage-threshold training data. We opted to compare BPNet-128 at a lower resolution (instead of base-resolution) to make a direct comparison across models, since the robustness metric is sensitive to bin-resolution. Indeed, models trained with augmentations yielded improved robustness compared to models without augmentations, especially when trained on peak-centered data (Fig. 3b). On the other hand, models that were trained on coverage-threshold data already benefited to a large degree on the non-centered, random nature of the epigenomic profiles. This could explain the observation that in BPNet-128 with RC augmentation alone was sufficient, although Basenji-128 still benefited from both augmentations. Surprisingly, we observed that models with similar prediction performance could yield large differences in their robustness levels, demonstrating that prediction performance and model robustness are not strictly correlated. This suggests that robustness can be utilized along with generalization performance as an additional metric to facilitate model selection.

### Comparing quantitative model architectures

The space of binary model architectures has been well explored; however, the exploration in quantitative models has been more limited. Existing quantitative models often have complex designs, including dilated convolutions with skipped connections and task-specific output-heads. It remains unclear to what extent that complex model structures are needed to fit quantitative data. We therefore wanted to address two questions: (1) could standard convolutional neural networks (CNNs) that were successful on binary classifications have similar success at quantitative regressions, and (2) could further exploration of the architectures improve performance?

To address these questions, we benchmarked a baseline CNN – with 3 convolutional layers and 1 fully connected hidden layer – at base and 32 bin-resolutions. We created 2 versions of each model where predictions are made based on task-specific output-heads, where each task is given a nonlinear prediction module or all predictions are based on a linear mapping from a single representation, i.e. the common multi-task approach. In addition, we set the first-layer activations of each model either to be rectified linear units (ReLU) or exponential activations, which has been shown to improve the quality of learned motif representations^32^. To test the benefits of a wider receptive field to give context to the patterns learned in lower layers, we created an augmented version of the baseline models by adding a residual block^33^, each with 4 dilated convolutional layers^34,35^, after each of the first two convolutional layers in a manner similar to ResidualBind^36^. Together, this results in a total of 8 custom models (see Methods for details).

Surprisingly, we found that baseline CNNs perform on par with Basenji- and BPNet-based models, with the exception of Basenji-32 (Table 1). This shows that simple model architectures can be effective at predicting epigenomic profiles as a quantitative regression. On the other hand, including dilated residual blocks, but with components arranged differently than Basenji, substantially improved performance at both tested resolutions for ResidualBind. Interestingly, task-specific output-heads consistently yielded better performance versus a single output-head, albeit the effect was variable and small. Moreover, exponential activations yielded comparable results to ReLU-based models, suggesting that high-divergence activations do not negatively affect the ability to make quantitative predictions. Together, this demonstrates that design considerations for quantitative models are largely under-explored and can greatly improve performance.

**Table 1.** Performance comparison across quantitative architectures. Table shows the whole-chromosome Pearson’s r (averaged across cell lines) for various quantitative models with different activations, output-heads, and trained on different target resolutions. For activations, Exponential refers to the application of it to only the first-layer filters while ReLU is used in deeper layers.

|  |  |  | Pearson's r (whole) |  |
| --- | --- | --- | --- | --- |
| Model | Activation | Output-head | Base | 32 Bin |
| BPNet | ReLU | Task-specific | 0.601 | 0.583 |
| Basenji | GELU | Single | 0.617 | 0.654 |
| CNN | ReLU | Single | 0.600 | 0.605 |
|  |  | Task-specific | 0.607 | 0.604 |
|  | Exponential | Single | 0.599 | 0.600 |
|  |  | Task-specific | 0.599 | 0.601 |
| ResidualBind | ReLU | Single | 0.642 | 0.655 |
|  |  | Task-specific | 0.637 | 0.670 |
|  | Exponential | Single | 0.655 | 0.665 |
|  |  | Task-specific | 0.654 | 0.677 |

### Benchmarking model performance across binary and quantitative models

Although quantitative models were developed with the aim of preserving more information about epigenomic profiles, directly comparing the different prediction formats between binary and quantitative tasks is not straightforward. To bridge this gap, we developed a way to directly compare binary models to quantitative models by converting predictions from one format to the other (Fig. 4a). To convert binary predictions to a quantitative format, we treated the logits (i.e. before the output activation function) as the predicted coverage values. While binary models are not trained to learn signal strength, the model’s confidence can be encoded in the unbound logits. Thus, binary models can now be evaluated with quantitative metrics. Moreover, to convert quantitative predictions to a binary format, we calculated the average coverage predictions at positive regions and negative regions based on corresponding binary-labelled data. These two distributions can be used to calculate standard classification metrics, such as the area-under the precision-recall curve (AUPR) and area-under the receiver-operating-characteristic curve (AUROC).

**Figure 4.**
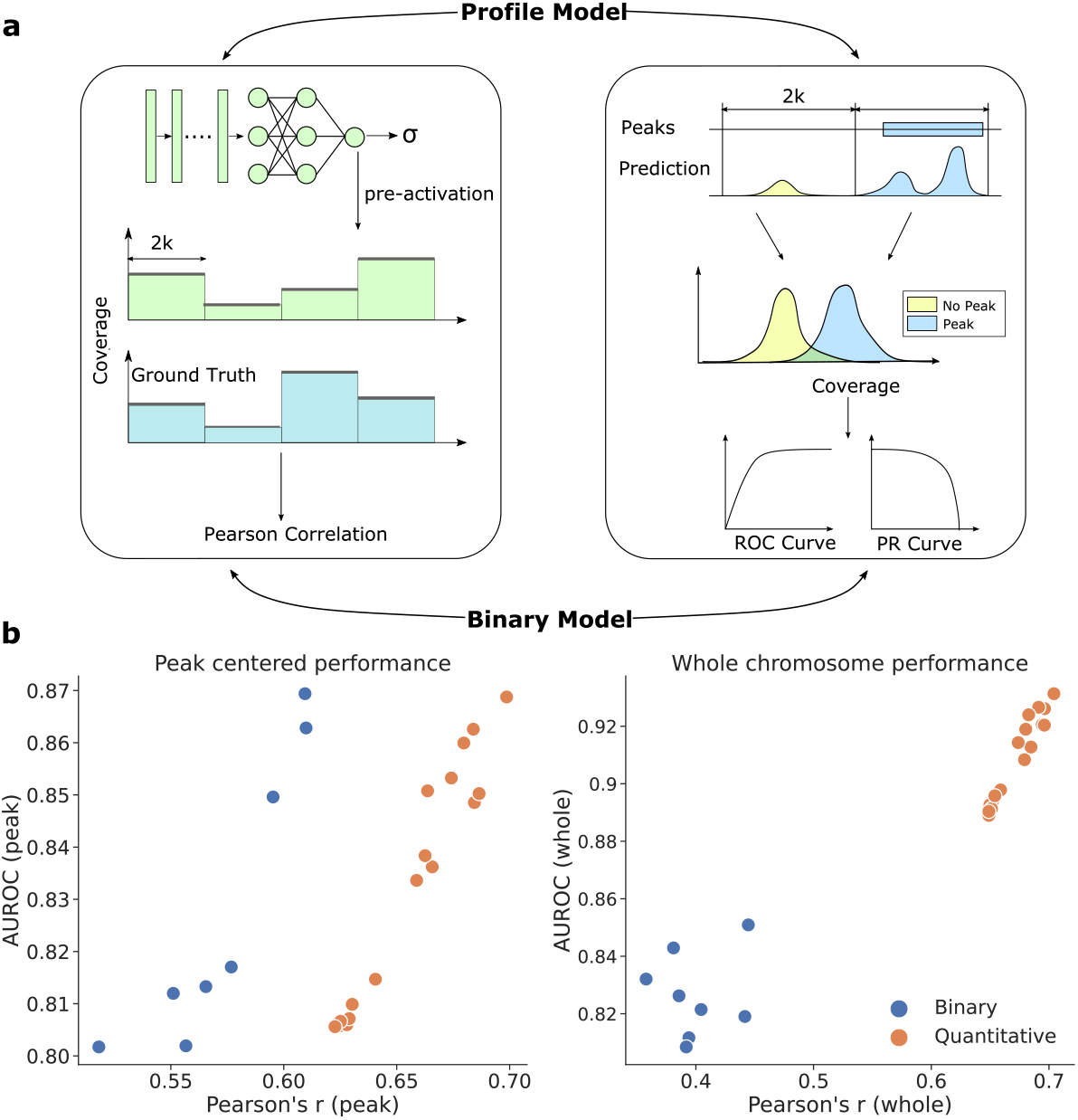
Performance comparison between binary and quantitative models. (**a**) Schematic overview of prediction task conversion. Binary models can use logits to generate continuous ‘coverage-like’ values to calculate regression metrics. On the other hand, the coverage predictions of quantitative models can be grouped according to binary labels (i.e. peak and no peak groups) to calculate standard classification metrics. (**b**) Scatter plot of the classification-based AUROC versus the regression-based Pearson’s r for various binary models (blue) and quantitative models (orange) on peak-centered test data (left) and whole-chromosome test data (right). Metrics represent the averaged value across cell lines.

Using this task-conversion evaluation framework, we directly compared the performance of various quantitative models with various binary models (Extended Data Table 2). Interestingly, when evaluating on peak-centered data, several binary models yielded similar (if not better) AUROC and AUPR compared to quantitative models (Fig. 4b and Extended Data Figure 3a). However, when converting the binary models to quantitative metrics, quantitative models outperformed all binary models. This effect became more pronounced when evaluation was extended across the whole chromosome, where all quantitative models yielded better performance across both metrics (Fig. 4c and Extended Data Figure 3b). Together, this demonstrates that while some binary models can be competitive with quantitative models within high-activity functional sites, quantitative models tend to yield better overall performance across whole chromosomes.

### Out-of-distribution generalization: variant effect prediction

A major downstream application for DL models that learn sequence-function relationships is to utilize them to score the functional effects of mutations. High-performing models can inform how their predictions change relative to wild-type when queried with a new mutated sequence. Thus, we benchmarked each model on this out-of-distribution (OOD) generalization task by validating predictions with experimental data from the CAGI5 Competition^37,38^, similar to previous studies^5,27^. CAGI5 dataset consists of massively parallel reporter assays (MPRAs) that measure the effect size of single-nucleotide variants through saturation mutagenesis of 15 different regulatory elements across different cell-types. Instead of a standard approach that makes a single prediction based on a sequence centered on the variant-of-interest, robust predictions were calculated by introducing random translations and averaging the central overlapping region, similar to our robustness test (see Methods). Robust predictions were calculated separately for reference and alternative alleles and the effect size was calculated based on their log2 fold change. An anecdotal visualization shows that the variant effect predictions by quantitative models are qualitatively effective despite being trained on OOD data – i.e. chromatin accessibility in different cell lines (Fig. 5a).

**Figure 5.**
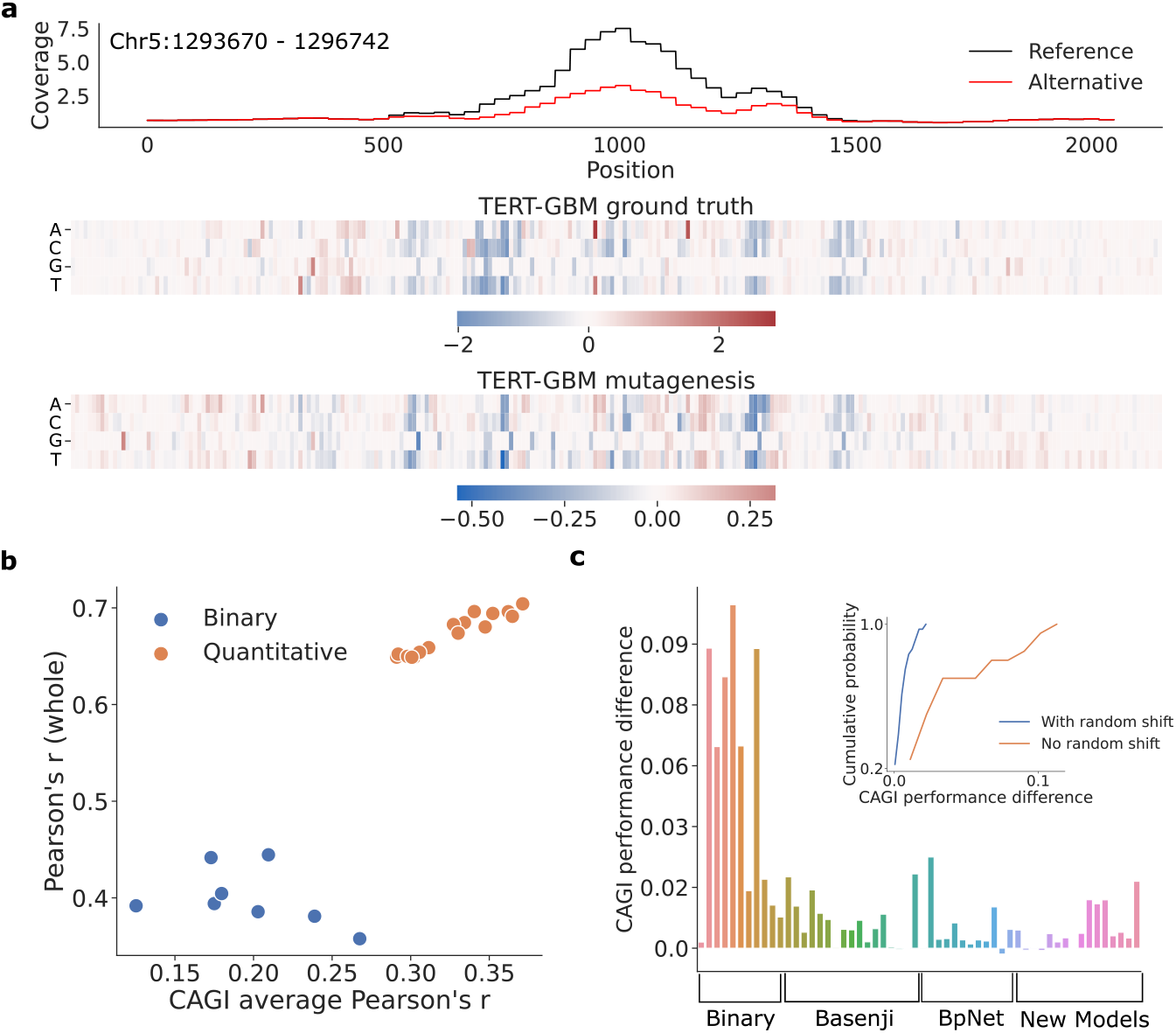
Comparison of functional effect predictions. (**a**) Example visualization of predictions of a sequence with a reference allele (black curve) and an alternative allele (red curve) for a given mutation. Below, heat maps show the experimental measurements of variant effects for the TERT promoter in a GBM cell line (ground truth) and the predicted variant effects from ResidualBind-32. (**b**) Scatter plot of the prediction performance across whole-chromosome test set (*y*-axis) and the average CAGI5 performance (*x*-axis). Each dot represents a different model. (**c**) Bar plot shows the CAGI5 performance difference between robust predictions minus standard predictions. Each bar represents a different model. Groups of models represent different training strategies or target resolutions. Inset shows the cumulative distribution of variant effect performance differences for models trained with and without random shift data augmentation.

By benchmarking various models, we found that quantitative models consistently outperformed binary models (Fig. 5b). In addition, by cross-comparison with prediction performance, we found that whole-chromosome generalization is a reliable metric for variant effect performance. As a control, we also compared whether robust predictions are beneficial for predicting variant effects compared to the standard approach of employing a single-pass prediction centered on the variant-site. Strikingly, we found that 50 out of 56 models performed better using robust predictions (Fig. 5c). Upon further investigation, models that did not employ random shift data augmentations were the ones that indeed benefited the most from robust predictions (Fig. 5c, inset). This suggests that robust models yield more trustworthy variant effect predictions, but our post hoc workaround of making predictions more robust could improve the efficacy of less robust models.

### Model interpretation

A major downstream application of genomic DL models is interpretability analysis, which can lead to the discovery of functional motifs and their complex interactions^39^. Here we compare binary and quantitative models across several common interpretability approaches: motif discovery through filter visualizations, foot-printing motifs at base-resolution using attribution methods, and quantitatively testing hypotheses *in silico* using global importance analysis (GIA).

#### Filter interpretability

First-layer convolutional filters provide an inductive bias to learn translational patterns such as motifs. However, the extent that they learn interpretable motifs largely depends on design choices, such as the max-pool size^40^, activation function^32^, or even the utilization of batch normalization^41^. However, it is not clear whether the same design principles established for binary models extends to quantitative models. To evaluate which models yield better motif representations, we visualized the first-layer filters of various models according to activation-based alignments^4,42^ and compared how well they match motifs in the JASPAR database^43^ using Tomtom^44^, a motif search comparison tool. Since the absolute number of hits can be misleading because of low-quality hits from partial motifs, we also consider the q-value, which specifies the confidence level of motif-filter matches. We found that among models that employ ReLU-based activations, binary models generally yield a lower hit-ratio as well as higher *q*-values, which indicates poor quality hits (Extended Data Table 2). However, models that employ exponential activations in the first convolutional layer yield higher hit-ratios and higher quality hits for both binary and quantitative models. We did not notice any significant differences in learning motif representations across architectures for a given activation. This suggests that the improved predictions do not necessitate learning better motif representations in the first layer, but the design principle using exponential activations can greatly improve the interpretability of learned motifs in first layer filters for both binary and quantitative models.

#### Embedding representations and attribution maps

To visualize structure in the data as seen through the lens of the model, we embedded the penultimate (or bottleneck) representations of test sequences for a given class using Uniform Manifold Approximation and Projection (UMAP)^45^. ResidualBind-32 with exponential activation yielded distinct UMAP structures and clustered data points with similar profile distributions in each cell line (Supplementary Fig. 2). By exploring different regions of the UMAP embeddings for the PC-3 cell line as an example, we found that ResidualBind-32 is largely encoding the magnitudes and locations of the read distributions (Extended Data Figure 4). We generated attribution maps (based on saliency analysis) from different regions and found that the ResidualBind-32 has learned many known regulatory motifs, such as AP-1, SP1, and GABPA, among others (Extended Data Figure 4). Interestingly, many accessible regions with high functional activity for PC-3 contained repeated clusters of AP-1, suggesting that our model considers AP-1 to be a critical motif for accessible sites for PC-3. We also observed an unknown motif (ATAAA) that flanked the AP-1 motif in many attribution maps potentially corresponding to a Forkhead family transcription factor binding site^43^. Many of these motifs were observed in attribution maps of other models, but due to the lack of ground truth, a quantitative comparison remains difficult.

#### Global importance analysis

While attribution maps can help to identify and footprint putative motifs, they cannot quantify the importance of motifs beyond individual nucleotides. GIA is an interpretability approach that enables direct testing of hypotheses gained from attribution analysis^36^. GIA computes the effect size, or global importance score, of hypothesis patterns that are embedded within a population of background sequence, where the other positions are effectively randomized. This approach essentially marginalizes out any confounding patterns within any individual sequence, revealing the global importance of only the embedded patterns on model predictions. Using GIA, we test various hypotheses of AP-1, ATAAA and GATA motifs.

First, we used GIA to explore how the two flanking nucleotides adjacent to the core motif on either side and the central nucleotide in AP-1 influences accessibility predictions in PC-3 cells. We compared the results from our high performing models within each prediction task, quantitative ResidualBind-32 and ResidualBind-binary, both with exponential activations. Strikingly, we find that flanking nucleotide combinations relative to the AP-1’s core binding site can drive predictions by a factor of 2 to 3 (Fig. 6a), similar to what was observed previously^46,47^. A position-weight-matrix-based approach^48^, which considers each position independently, would score many AP-1 binding sites the same, despite their wide spread in functional activity. A similar observation was made for other models, both quantitative and binary (Extended Data Figure 5). This demonstrates that DL models consider complex higher-order dependencies of flanking nucleotides to be an important feature of TF binding sites, a well-known phenomenon^49,50^. Moreover, both binary and quantitative models can capture this information *de novo* from being trained on just the sequences.

**Figure 6.**
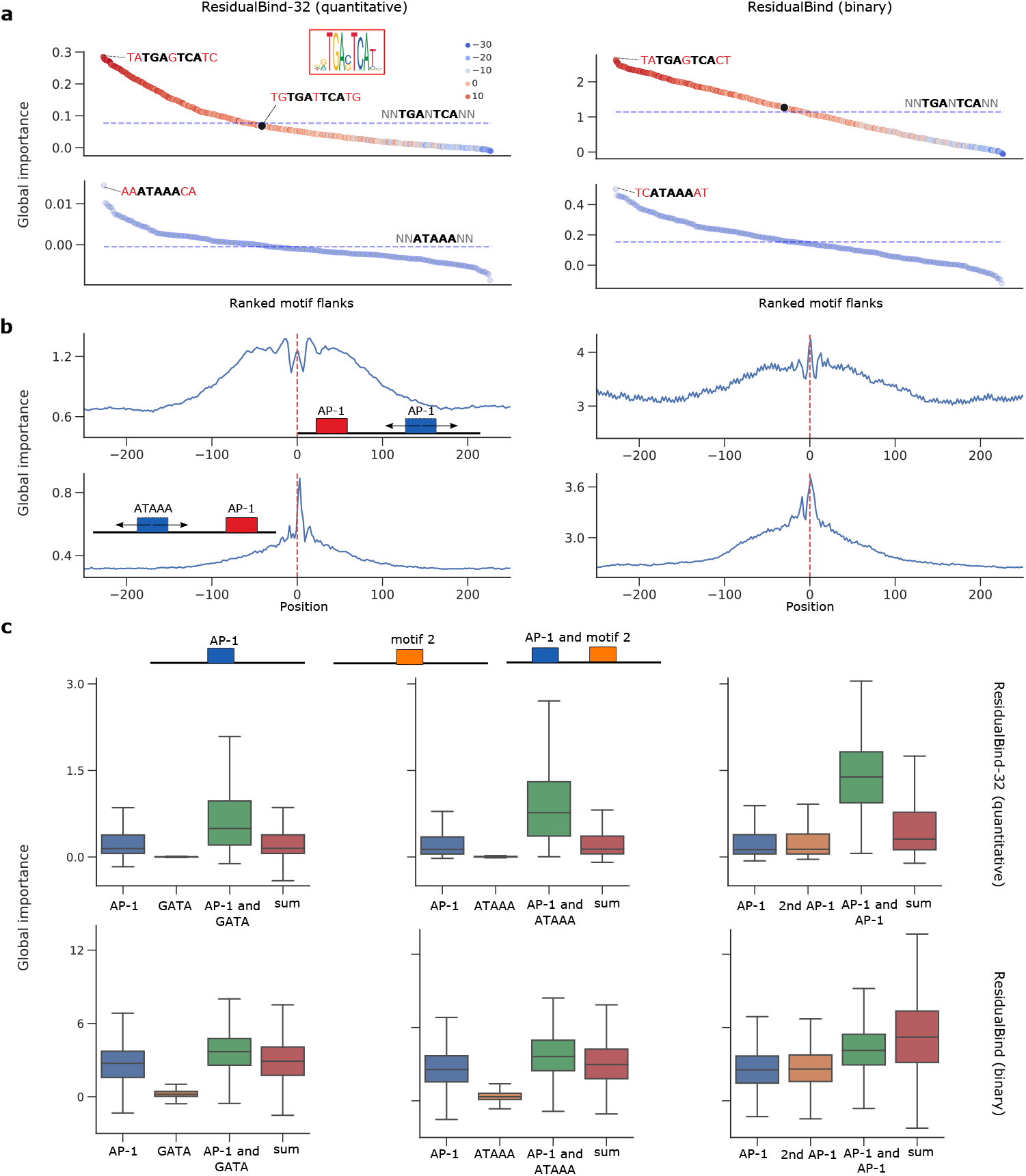
Interpretability analysis for ResidualBind-32 on PC-3 cell line. (**a**) GIA for optimal flanking nucleotides. Ranked plot of the global importance for each tested flank. Dashed line represents the global importance of the core motif with random flanks. The hue represents the position-weight-matrix score for an AP-1 motif from the JASPAR database (ID: MA0491.1). The black dot indicates ‘TGTGATTCATG’, which has a high position-weight-matrix score but yields a global importance close to the core motif with randomized flanks. **(b)** GIA for distance dependence between two motifs with optimized flanks. Global importance plot for sequences with an AP-1 motif fixed at the center of the sequence and another motif that is systematically placed in different locations. Positive and negative values represent the first positions the motifs were embedded to be non-overlapping. **(c)** GIA for cooperative interactions between AP-1 and another motif. Each subplot shows a box-plot of the global importance when each motif is placed in random background sequences individually and in combinations. For comparison, the sum of the global importance scores for each individual motif is also shown (sum). Box plots show the first and third quartiles, central line is the median, and the whiskers show the range of data with outliers removed.

We then explored to what extent the distance between motifs plays a role in model predictions. Specifically, we performed GIA experiments where the AP-1 motif was fixed at the center of the sequences and the position of the other motif (i.e. AP-1 or ATAAA) was varied (Fig. 6b). Interestingly, we found that two AP-1 motifs yielded a symmetric 50 bp window where predictions are plateaued, beyond which, the global importance begins to drop off for both models. On the other hand, the ATAAA motif exhibits an asymmetric distance dependence with a favorable location on the 3’ end flanking the AP-1 motif with a few nucleotide gap, beyond which there is a precipitous drop in global importance. This was also observed across other models, with variable magnitudes in effect size but a similar relative trend (Extended Data Figure 6). These results suggest that flanking nucleotides and distance dependence was consistently learned across quantitative and binary models.

From the attribution maps, it appears that multiple motifs are often present in combinations across accessible sites. To test whether ResidualBind-32 and ResidualBind binary have learned cooperativity between AP-1 and other motifs, we compared the global importance for the motifs embedded in sequences alone and in combinations with other motifs (at the optimal distance identified through the distance dependence GIA experiments). In ResidualBind-32, we observed the sum of individual effects were lower than when both motifs were present, indicating the model has indeed learned cooperative interactions (Fig. 6c). The effect size was varied across transcription factors, with a smaller effect observed for GATA∷AP-1 compared to other motifs, such as ATAAA∷AP-1 and AP-1∷AP-1. This suggests that cooperative interactions are strongly associated with chromatin accessibility levels. Surprisingly, a discrepancy arose for binary models, including in ResidualBind binary, for which there was no strong evidence that cooperativity was learned. These trends were also observed across other binary and quantitative models (Extended Data Figure 7).

Instead of directly imposing patterns on background sequences, we also conducted occlusion-based interventional experiments where we identified exact instances of the core motif for AP-1 and replaced them with randomized sequences across the test set – a global importance of motif occlusion within its natural sequence context. We find that the number of AP-1 motifs indeed drives high functional activity for PC-3, while other cell lines depend on a different distribution of motifs for their functional activity (Supplementary Fig. 3).

Together, the interpretations of quantitative models appear to be more consistent with each other than binary models. Despite under-performing on generalization tasks, well-trained binary models can largely capture similar biological interpretations as quantitative models, with the exception of cooperative interactions.

## Discussion

The variety of deep learning models being proposed to predict regulatory genomic tasks has increased substantially in recent years. The variations of proposed models, how the prediction tasks are framed, the composition of the data sets, and the tricks used for training make it challenging to assess which innovations are driving performance gains. Moreover, while many methods provide software to deploy their methods, which only includes their specific pipeline, it is often challenging to mix-and-match modeling innovations across methods. To address this gap, we introduced GOPHER to provide an evaluation framework to compare various modeling choices and enable a comprehensive and fair evaluation of existing and emerging DL models in regulatory genomics. While previous software, such as Janggu^51^ and Selene^52^, help to process biological data (mainly focused on peak-centered data) and high-level APIs to train neural network models in TensorFlow^53^ and Pytorch^54^, respectively, they do not focus on downstream evaluation across different prediction tasks. By contrast, GOPHER provides a comprehensive model evaluation framework that also supports data processing of peak-based binary classification and quantitative regression analysis of bigWig tracks, in addition to training custom deep learning models with various data augmentations. GOPHER also incorporates many popular model interpretability tools, such as first-layer filter visualization, global importance analysis, and attribution methods, including *in silico* mutagenesis^55,56^, saliency maps^57^, integrated gradients^58^, and SmoothGrad^59^.

Using GOPHER, we addressed several open questions: (1) how to fairly compare binary models and quantitative models; (2) how choice of loss function affects performance; (3) how dataset selection influences model performance; (4) how to compare quantitative models that making predictions at different resolutions; (5) how augmentation strategies influence model performance and robustness to translational perturbations; how modeling choices influence downstream (6) functional variant effect predictions and (7) model interpretability.

red While the study here focuses on ATAC-seq data, the specific claims of optimal architectures and training procedures may be nuanced across other data types, such as ChIP-nexus^26^ and CAGE-seq^60^. In such cases, additional considerations may arise, such as GC-bias and signal normalization. These were not investigated in this study.

Moreover, many of the explored architectures in this study relied on pure convolutional networks. The emergence of transformers^61^ could have better inductive bias to capture distal interactions, though the rationale for the benefits of convolution-based networks and transformer-based networks remains an ongoing research topic^62^. Due to the lack of established transformer-based models beyond Enformer^27^, we elected to focus only on convolution-based models in this study.

In addition, BPNet and Basenji were initially developed for predictions on very different data types and dataset sizes. Thus modifications had to be made to each model architecture to adapt it to the ATAC-seq data used in this study (see Methods for differences). These choices may have affected their performance. The fact that both models performed well suggests that our hyperparameter optimization mitigated any substantial disparities.

In general, our work largely supports that quantitative modeling yields better generalization (on average), both on held-out data and OOD variant effect predictions. Of course, well-tuned binary models can perform comparable to (or even better than) a poorly designed quantitative model. It remains unclear whether binary models are fundamentally limited based on their treatment of functional activity or whether incorporating more inactive regions during training would boost performance. Moreover, it is not clear whether the performance gains of quantitative models are due to learning better biological signals or whether they are just better at learning noise sources within sequencing experiments. One major limitation arises as a consequence of focusing on performance – treating experimental measurements as ground truth, despite biological variability across replicates and technical noise (eg. Supplementary Table 2). Thus, focusing on important downstream tasks, such as variant effect prediction and model interpretability, as was done here, provides a path to move beyond performance benchmarks to the beneficial use case of genomic DL models – biological discovery.

## Methods

### Training data

ATAC-seq (Assay for Transposase-Accessible Chromatin with high-throughput sequencing^63^) data for human cell lines were acquired from the ENCODE database^64^ – fold change over control bigWig files for quantitative analysis and IDR peak bed files for binary analysis – using experimental accessions in Supplementary Table 3. The bigWig tracks were log-fold-normalized for sequencing depth using control tracks as per the ENCODE data processing pipeline; no further processing was done. Each of the 15 cell lines were sub-selected based on a lower cross-correlation of coverage values at IDR peaks across cell lines below 0.75. Data from replicate 1 for each experiment was used to generate the train, validation, and test sets. Data from replicate 2 was used to assess the experimental ceiling of prediction performance.

#### Coverage-threshold data

Each chromosome is split into equal, non-overlapping input size chunks of 3 kb and each chunk is included in the dataset if the max coverage value for any of the targets is above the threshold. By default, coverage-threshold data employed a threshold of 2, unless specified otherwise. Each sequence that passed this threshold was included as part of the dataset and down-sampled to 2 kb with a strategy that depends on data augmentations (see below). The targets were then binned with non-overlapping windows according to the specified target resolution and was calculated online during training and testing. For any given coverage value array of length *L* and bin size *B*, it was reshaped into an array with shape (*B*, *L*/*B*) – down-sampling was achieved according to the mean within each bin.

#### Peak-centered data

For peak-centered datasets, we selected IDR bedfiles from the ENCODE experiments corresponding to the same replicate as the coverage-threshold data. The bed files of each cell line were merged into a single bed file, in a manner similar to Kelley et al^4^. The Basset data processing pipeline divides the genome into segments of length specified as the input size and merges peaks according to an overlap size parameter. Each sequence in the dataset contains at least one peak across all cell lines. Sequences containing an IDR peak for the cell line is given label ‘1’ otherwise label ‘0’.

#### Data splits

We split the dataset into training, validation, and test sets using chromosome 8 for test, chromosome 9 as validation and the rest as training (excluding Y chromosome). We also removed the unmappable regions across all data splits. The same split was applied to datasets to allow a direct comparisons across experiments.

### Held-out test evaluation

Pearson correlation can be calculated using the concatenated whole chromosome per cell line, which is referred to as Pearson’s r (whole), or per sequence correlation averaged across the test set when specified. The difference between these metrics manifests as a different mean in the correlation calculation; a global mean for whole chromosome versus a per sequence mean. Whole chromosome evaluation is calculated by concatenating the predictions for the entire chromosome 8 with the exception of unmappable regions. A per sequence Pearson correlation was calculated for peak-centered data, test selection analysis, and robustness analysis, unless specified otherwise. For a compilation of all model evaluations see Supplementary Data 1.

#### Scaling predictions

Predictions were scaled to address the large discrepancy between predictions and experimental values for shape-based loss functions (eg. Pearson’s r). Though we found that applying it to other losses also yielded slightly better performance. This was accomplished by calculating a global scaling factor per cell line, which is computed as the ratio of the mean of experimental and predicted coverage values across the entire test chromosome, and multiplying the scaling factor to the predictions.

### Models

#### Basenji

Basenji-based model is composed of a convolutional block, max-pooling with pool size of 2 (which shrinks the representations to 1024), 11 residual blocks with dilated convolutional layers, followed by a final convolutional layer. The convolutional block consists of: GELU activation^65^, convolutional layer with a kernel of width 15, batch normalization^66^. The residual block is composed of: GELU activation, dilated convolutional layer with a kernel of width 3 and half the number of filters and a dilation rate that grows by a factor of 1.5 in each subsequent residual block, batch normalization, GELU activation, convolutional layer with width 1 and the original number of filters, batch normalization, and dropout with a rate 0.3. Each residual block has a skipped connection to the inputs to residual block. An average pooling layer is applied to the final output convolutional layer to shrink the representations to the corresponding target resolution. A dense layer with softplus activations following the last convolutional block then outputs the predictions target. In case of base resolution, the first max-pool size is set to 1.

The original Basenji model employs multiple convolutional blocks with a max pooling of size 2 to reduce the dimensions of the sequence to 1024 units, upon which 11 residual blocks are applied. Since our input size is 2048 bps, we employed a single convolutional block to achieve the same dimensions as the original Basenji model. We performed hyperparameter search of the number of convolutional filters in each layer to optimize for the ATAC-seq dataset used in this study. For additional details of specific hyperparameters, see Supplementary Data 2.

#### BPNet

BPNet consists of a convolutional layer, followed by 9 dilated convolutional layers with progressively increasing dilation rates (scaled by powers of 2) that each have a residual connection to the previous layer. Task-specific output-heads, each with a separate transpose convolution, is built upon the final residual layer. To adapt BPNet to lower resolutions, all predictions are initially made at base-resolution followed by an average pooling layer for each task-specific output-head, with a window size and stride that matches the target resolution.

A key difference with the original BPNet architecture is that the negative strand, bias track and read counts output-head was not used throughout this study. Moreover, the original loss function was not employed here as we found better success with the modified BPNet using a Poisson NLL loss. This may be attributed to the lower resolution in read coverage for bulk ATAC-seq, or due to original model targeting raw read count instead of fold change over control tracks, though further investigation is needed to understand the disparity. These modifications may have affected the performance of BPNet. We optimized hyperparameters of the model, focusing on the number of filters in each layer and the kernel size of the transpose convolution in the task-specific output heads (Supplementary Fig. 1). The specific choices of hyperparameters in BPNet can be found in Supplementary Data 2.

#### CNN-baseline

The CNN baseline model is composed of 3 convolutional blocks, which consist of a 1D convolution, batch normalization, activation, max pooling and dropout, followed by 2 fully-connected blocks, which includes a dense layer, batch normalization, activation, and dropout. The first fully connected block scales down the size of the representation, serving as a bottleneck layer. The second fully-connected block rescales the bottleneck to the target resolution. This is then reshaped to match the number of bins × 8. For instance, the number of hidden units for models at 32 bin target resolution are 2048/32 = 64 × 8, then reshaped to (64, 8). Base resolution models set the hidden units to 2048 × 8 then reshaped to (2048, 8). This is followed by another convolutional block. The representations from the outputs of the convolutional block is then input into task-specific output heads or is directly fed to a linear output layer with softplus activations. For task-specific output heads, each head consists of a convolutional block followed by a linear output layer with softplus activations. The activation of the first layer is either exponential or ReLU, while the rest of the hidden layer activations are ReLU. The specific hyperparameters of each layer, including the dropout rates, are specified in detail in Supplementary Data 2.

#### ResidualBind-base

ResidualBind-base builds upon the CNN-baseline models by adding a residual block after the first 3 convolutional layers. The first two residual blocks consist of 5 dilated convolutional layers and the third residual block consists of 4 dilated convolutional layers. Similar to CNN-baseline models, this is then followed by 2 fully connected blocks, which are reshaped to a shape (2048, 8), and a convolutional block. Here, another residual block that consists of 5 dilated convolutional layers was applied. This is then fed into an output head, which has the same composition as the CNN-baseline. The details of model architecture and hyperparameters can be found in Supplementary Data 2.

#### ResidualBind-32

ResidualBind-32 also builds upon the CNN-baseline models by adding a residual block after the first 3 convolutional layers, but with a few key differences from ResidualBind-base. The third residual block consists of 3 dilated convolutional layers instead of 4. Moreover, ResidualBind-32 does not go through a bottleneck layer that is prototypical of the CNN-baseline design. For task-specific output heads, the representations of the third residual block are input into a convolutional block followed by a task-specific output heads similar to the CNN-baseline models. For a single output head, the representations of the third residual block are input into a position-wise fully connected block followed by a linear output layer. The details of model architecture and hyperparameters can be found in Supplementary Data 2.

#### Binary models

Four main model structures are used for binary models. One fine-tuned Basset^4^ structure and three re-purposed quantitative models structures: Basenji, CNN-base, and ResidualBind-base. Basset is composed of three blocks of convolutional layer followed by batch normalization, activation and max-pooling. The output is then flattened and fed into 2 fully connected layers with dropout and an output layer with sigmoid activations. Basset hyperparameters were optimized for the binary version of the ATAC-seq dataset in a similar manner to Basenji. For Basenji-binary, CNN-binary and ResidualBind-binary, their structure highly resembles the quantitative model based on a single output-head. For CNN-binary and ResidualBind-binary, we apply a fully connected output layer with sigmoid activations to the bottleneck layer. For Basenji-binary, we take the penultimate representation and perform a global average pool, followed by a fully connected output layer. The details of model architecture and hyperparameters can be found in Supplementary Data 2.

#### Training

Each quantitative model was trained for a maximum of 100 epochs using ADAM^67^ with default parameters. Early stopping was employed with a patience set to 6 epochs (monitoring validation loss as a criterion). By default, models were trained with random reverse-complement and random shift data augmentations unless specified otherwise.

Quantitative CNN and ResidualBind (base and 32 bin-resolution), along with binary versions of these models, were trained for a maximum of 100 epochs using ADAM with default parameters. Early stopping with a patience of 10 was used. The initial learning rate was set to 0.001 and decayed by a factor of 0.2 when the loss function did not improve after a patience of 3 epochs.

### Data augmentations

#### Random shift

Random shift is a data augmentation that randomly translates the input sequence (and corresponding targets) online during training. All datasets were generated with input size set to 3,072 bp. When random shift is used, for each mini-batch, a random sub-sequence of 2,048 bp and its corresponding target profile was selected separately for each sequence. When random shift is not used, the central 2,048 bp is selected for all sequences in the mini-batch.

#### Reverse-complement

Reverse-complement data augmentation is employed online during training. During each mini-batch, half of training sequences were randomly selected and replaced by their reverse-complement sequence. For those sequences that were selected, the training target was correspondingly replaced by the reverse of original coverage distribution.

### Hyperparameter search

The ATAC-seq datasets in this study differ greatly in complexity, i.e. size and coverage distribution, from the original Basenji and BPNet studies. Therefore, we performed a hyperparameter search for each base architecture for our ATAC-seq dataset (Supplementary Fig.1). We used WandB^68^ to keep track of the model choices and for visualization. We fine-tuned Basenji and BPNet at 128 bp and base resolution, respectively, which represent the original resolutions for these models. We also kept their original training set selection strategy, that is, we trained Basenji on coverage-threshold data and BPNet on peak-centered data. For Basenji, the number filters across the convolutional layers were varied as well as the presence or absence of dropout layers (fixed rate for each layer). For BPNet, we performed a hyperparameter search over the number of convolutional filters in each layer and the kernel size in the task-specific output heads. We employed the original data augmentations (i.e. random reverse-complement and random shifts for Basenji and only random reverse-complement for BPNet). For each model, we used the Poisson NLL loss function. We originally used a MSE and multinomial NLL loss for BPNet, but found that optimization using Poisson NLL yielded better performance. The models were trained for maximum of 40 epochs with an Adam optimizer^67^ using default parameters. Initialization was given according to Ref.^33^. The optimal set of hyperparameters for each model was selected based on the lowest validation loss and the final architectures are given in Supplementary Data 2.

### Robustness test

To measure the robustness to translational perturbations, we analyzed the sequences within the held-out test chromosome that were identified to contain a statistically significant peak for the given cell-type under investigation. This ensures that the robust predictions are only considered for genomic regions that exhibit statistically significant coverage values. Specifically, we took each 3072 bp sequence in the dataset and generated 20 contiguous sub-sequences of length 2048 bp. Each sub-sequence was sent through the model to get a prediction, and all of the predictions were aligned based on the sub-sequence. All sub-sequences contain a center 1024 bp window that overlaps. Standard deviation is calculated for each position across these 20 sequences and averaged across the length of prediction. The average sequence coverage across 20 sequences was used to normalize the average standard deviation to make it invariant to scale. Therefore variation score for each sequence is calculated as average per position standard deviation divided by average sequence coverage. A higher variation score corresponds to a less robust model, while a lower variation score corresponds to more stable predictions, irrespective of translations to the inputs. Due to binning artifacts, we only compare this robustness test for models that share the same bin-resolution.

### Variant effect prediction

#### Dataset

The CAGI5 challenge dataset was used to benchmark model performance on variant calling. Each regulatory element ranges from 187bp - 600bp in length. We extracted 2048 bp sequences from the reference genome centered on each regulatory region of interest and converted it into a one-hot representation. Alternative alleles are then substituted correspondingly to construct the CAGI test sequences.

#### Standard predictions

For a given model, the prediction of 2 sequences, one with a centered reference allele and the other with an alternative allele in lieu of the reference allele, is made and the coverage values are summed separately for each cell-type. For each sequence, this provides a single value for each cell-type. The cell-type agnostic approach employed in this study then uses the mean across these values to calculate a single coverage value. The effect size is then calculated with the log-ratio of this single coverage value for the alternative allele and reference allele, according to: log(alternative coverage / reference coverage).

#### Robust predictions

For a given model, robust predictions were made by: 1) sampling 20 randomly shifted sequences centered on a variant-of-interest, 2) sending them through the model to get coverage predictions for each cell-type, 3) align predictions based on the shifted sub-sequences, 4) calculating the mean coverage within overlapping 1024 bp region for each cell-type, and 5) averaging the mean coverage values across cell-type. This was done separately for the reference allele and the alternative allele, and the effect size was calculated similar to the standard predictions as the log-ratio.

#### Evaluation

To evaluate the variant effect prediction performance, Pearson correlation was calculated within each CAGI5 experiment between the experimentally measured and predicted effect size. The average of the Pearson correlation across all 15 experiments represents the overall performance of the model. A full list of variant effect prediction performances for models can be found in Supplementary Data 3.

### Model interpretability

#### Tomtom

The motif comparison tool Tomtom^44^ was used to match the position probability matrix of the first convolutional layer filters (calculated via activation-based alignments^42^) to the 2022 JASPAR nonredundant vertebrates database^43^. Matrix profiles MA1929.1 and MA0615.1 were excluded from filter matching to remove poor quality hits; low information content filters then to have a high hit rate with these two matrix profiles. Hit ratio is calculated by measuring how many filters were matched to at least one JASPAR motif. Average q-value is calculated by taking the average of the smallest q-value for each filter among its matches.

#### Attribution analysis

Attribution analysis was based on grad-times-input with saliency maps^57^. For a given model, gradients of the prediction with respect to a given cell-type were calculated with respect to the input sequence to yield a *L × A* map, where *L* is the length of the sequence and *A* is 4 – one for each nucleotide. Each saliency map was multiplied by the input sequence, which is one-hot, to obtain just the sensitivity of the observed nucleotide at each position. A sequence logo was generated from this by scaling the heights of the observed nucleotide, using Logomaker^69^.

#### Global importance analysis

For global importance analysis^36^, we generated background sequences by performing a dinucleotide shuffle of 1,000 randomly sampled sequences from those within our coverage-threshold test set. The global importance is calculated via the average difference in predictions of background sequences with embedded patterns-under-investigation and without any embedded patterns. For quantitative models, the predictions represent the average coverage predictions for the cell-type under investigation. For binary models, the predictions represent the logits for the cell-type under investigation.

##### GIA for flanking nucleotides

We fixed the core motif at the center of all background sequences, i.e. starting at position 1024, and varied the 2 flanking nucleotides on each side (and the central nucleotide for only AP-1) by separately performing a GIA experiment for all possible combinations of flanking nucleotides.

##### GIA for distance-dependent motif interactions

To quantify the functional dependence of the distance between 2 motifs with optimized flanks, we fixed the position of 1 motif at the center of the sequence, i.e. starting at position 1024, and then systematically performed a GIA experiment with the second motif at different locations ensuring no overlap. This experiment provides a global importance score for the 2 motifs at different distances in both positive and negative directions.

##### GIA for motif cooperativity

To quantify whether motifs are cooperatively interacting, we inserted each motif (with optimized flanks) at the corresponding position (1024 for motif 1 and best position for interaction for motif 2 based on the distance-dependent GIA experiments) individually and in combinations. We then compared the global importance when both motifs are embedded in the same sequence versus the sum of the global importance when only one motif is embedded.

#### Occlusion-based experiments

We randomly sampled 10,000 sequences from those within our coverage-threshold test set. We performed a string search looking for exact matches to the core motif of AP-1, i.e. TGA-TCA, where the - can be any nucleotide. For each cell-type, we grouped the sequences according to the number of instances that the core AP-1 motif was observed – 1 observed motif, 2 observed motifs, and 3 or more observed motifs. For each group, we replaced the core motif with randomized sequences. Due to spurious patterns from randomized sequences, we performed a GIA experiment where 25 randomized sequences were embedded in lieu of the core binding site and the model predictions were averaged – first across the coverage for the cell-type under investigation, then across the 25 randomized sequences. This effectively marginalizes out the impact of the motif for a given sequence. This occlusion-based (or conditional) GIA experiment was done for each sequence in each group.

## Data availability

The processed ATAC-seq data that support the findings of this study are available in Zenodo, https://doi.org/10.5281/zenodo.6464031.

## Code availability

The code to reproduce results and figures in this study is available at Zenodo, https://doi.org/10.5281/zenodo.6464031. The open-source project repository is available at GitHub, https://github.com/shtoneyan/gopher. Stable version of code used for generating results in paper is available through Zenodo, https://doi.org/10.5281/zenodo.6977214

## Acknowledgements

This work was supported in part by funding from the Simons Center for Quantitative Biology at Cold Spring Harbor Laboratory. This work was performed with assistance from the US National Institutes of Health Grant S10OD028632-01. The authors would like to thank Jakub Kaczmarzyk and other members of the Koo lab for helpful discussions.

## Author contributions statement

ST, ZT, and PKK conceived of the experiments. ST and ZT wrote the code base, conducted the experiments, analyzed and interpreted the results. All authors contributed to the manuscript.

