## Extended Data for "Evaluating deep learning for predicting epigenomic profiles"

**Extended Data Table 1.** Comparison of models trained on peak-centered versus coverage-threshold data. Basenji-128 and BPNet-base with and without data augmentations (i.e. random RC and translational shift) were evaluated according to the mean-squared error (MSE) and Pearson's r on different held-out test sets, i.e. peak-centered or whole chromosome. Red values highlight the better metric when comparing different training sets for a given model and data augmentation.

| Model | Data augmentation | Training data | Test set selection | | | |
|---|---|---|---|---|---|---|
| | | | MSE Peak | MSE Chrom | Pearson's r Peak | Pearson's r Chrom |
| Basenji-128 | None | Peak-centered | 2.06 ±0.020 | 0.664 ±0.020 | 0.582 ±0.003 | 0.413 ±0.003 |
| | | Thresholded | 2.21 ±0.059 | 0.580 ±0.007 | 0.590 ±0.003 | 0.456 ±0.001 |
| | RC and shift | Peak-centered | 1.78 ±0.032 | 0.576 ±0.007 | 0.604 ±0.002 | 0.441 ±0.001 |
| | | Thresholded | 1.89 ±0.018 | 0.545 ±0.002 | 0.610 ±0.001 | 0.464 ±0.000 |
| BPNet-base | None | Peak-centered | 2.22 ±0.003 | 0.882 ±0.004 | 0.550 ±0.001 | 0.374 ±0.00130 |
| | | Thresholded | 2.48 ±0.019 | 0.683 ±0.002 | 0.545 ±0.000510 | 0.422 ±0.001 |
| | RC and shift | Peak-centered | 2.06 ±0.028 | 0.770 ±0.018 | 0.565 ±0.002 | 0.401 ±0.00225 |
| | | Thresholded | 2.29 ±0.013 | 0.640 ±0.003 | 0.563 ±0.00104 | 0.429 ±0.002 |

**Extended Data Table 2.** Performance comparison across prediction tasks. Various models – trained on binary and coverage data, different output heads, and activation functions – were compared across prediction tasks within whole-chromosome data (whole) and peak-centered data (peak) and filter analysis – hit-ratio of the filters to the JASPAR database and the average of the best q-value for each filter.

| Model | Resolution | Output Head | Activation | chrom AUROC | chrom AUPR | chr Pearson R | peak AUROC | peak AUPR | peak Pearson R | hit ratio | q value |
|---|---|---|---|---|---|---|---|---|---|---|---|
| Binary Models | | | | | | | | | | | |
| Basenji | Binary | Single | GELU | 0.819 | 0.307 | 0.442 | 0.802 | 0.576 | 0.518 | 0.527 | 0.204 |
| | | | Exponential | 0.851 | 0.292 | 0.445 | 0.817 | 0.607 | 0.577 | 0.676 | 0.153 |
| Basset | Binary | Single | ReLU | 0.812 | 0.314 | 0.394 | 0.802 | 0.592 | 0.557 | 0.503 | 0.199 |
| | | | Exponential | 0.821 | 0.314 | 0.404 | 0.812 | 0.603 | 0.551 | 0.833 | 0.104 |
| CNN | Binary | Single | ReLU | 0.808 | 0.305 | 0.392 | 0.813 | 0.608 | 0.565 | 0.682 | 0.106 |
| | | | Exponential | 0.826 | 0.334 | 0.386 | 0.850 | 0.657 | 0.595 | 0.859 | 0.064 |
| Residual | Binary | Single | ReLU | 0.843 | 0.361 | 0.381 | 0.863 | 0.677 | 0.610 | 0.448 | 0.178 |
| | | | Exponential | 0.832 | 0.346 | 0.358 | 0.869 | 0.688 | 0.609 | 0.833 | 0.074 |
| Quantitative Models | | | | | | | | | | | |
| CNN | 1 | Task-Specific | ReLU | 0.898 | 0.478 | 0.659 | 0.815 | 0.636 | 0.641 | 0.823 | 0.074 |
| | | | Exponential | 0.889 | 0.470 | 0.649 | 0.806 | 0.629 | 0.624 | 0.859 | 0.050 |
| | | Single | ReLU | 0.894 | 0.472 | 0.652 | 0.806 | 0.625 | 0.625 | 0.797 | 0.070 |
| | | | Exponential | 0.893 | 0.469 | 0.650 | 0.806 | 0.627 | 0.628 | 0.844 | 0.062 |
| | 32 | Task-Specific | ReLU | 0.891 | 0.470 | 0.651 | 0.807 | 0.628 | 0.629 | 0.812 | 0.072 |
| | | | Exponential | 0.889 | 0.468 | 0.649 | 0.807 | 0.629 | 0.625 | 0.828 | 0.072 |
| | | Single | ReLU | 0.896 | 0.474 | 0.654 | 0.810 | 0.629 | 0.630 | 0.812 | 0.081 |
| | | | Exponential | 0.890 | 0.469 | 0.649 | 0.806 | 0.626 | 0.623 | 0.839 | 0.050 |
| Residual | 1 | Task-Specific | ReLU | 0.908 | 0.520 | 0.679 | 0.834 | 0.667 | 0.659 | 0.443 | 0.173 |
| | | | Exponential | 0.921 | 0.538 | 0.694 | 0.849 | 0.680 | 0.684 | 0.823 | 0.070 |
| | | Single | ReLU | 0.913 | 0.524 | 0.685 | 0.836 | 0.666 | 0.666 | 0.505 | 0.179 |
| | | | Exponential | 0.920 | 0.540 | 0.696 | 0.850 | 0.683 | 0.686 | 0.833 | 0.067 |
| | 32 | Task-Specific | ReLU | 0.926 | 0.552 | 0.696 | 0.863 | 0.699 | 0.684 | 0.496 | 0.175 |
| | | | Exponential | 0.931 | 0.561 | 0.704 | 0.869 | 0.707 | 0.699 | 0.867 | 0.063 |
| | | Single | ReLU | 0.919 | 0.531 | 0.680 | 0.851 | 0.681 | 0.664 | 0.516 | 0.194 |
| | | | Exponential | 0.927 | 0.544 | 0.691 | 0.860 | 0.691 | 0.680 | 0.852 | 0.068 |
| Basenji | 128 | Single | GELU | 0.924 | 0.532 | 0.683 | 0.853 | 0.680 | 0.674 | 0.531 | 0.178 |
| BpNet | 1 | Task-Specific | ReLU | 0.914 | 0.507 | 0.674 | 0.838 | 0.660 | 0.663 | 0.395 | 0.080 |

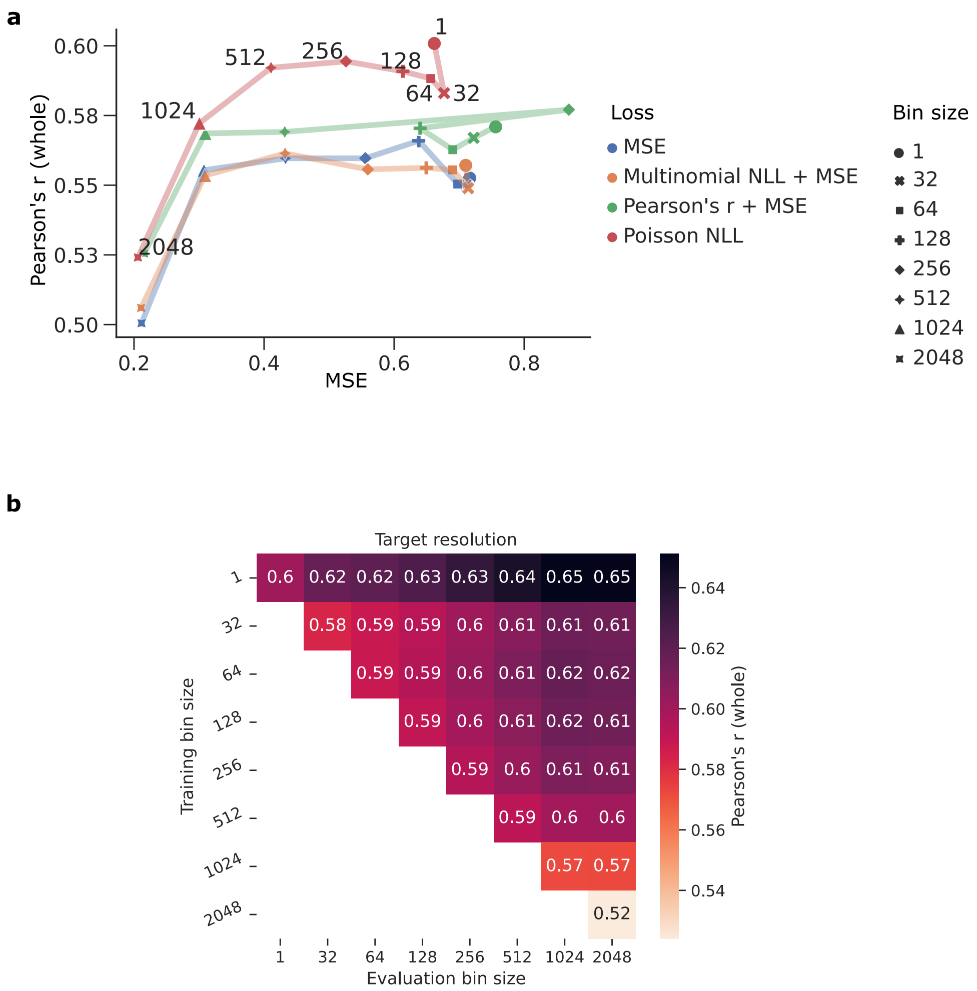


**Extended Data Figure 1.** Evaluation of BPNet-based quantitative models. (**a**) *Loss function analysis.* Scatter plot of the whole-chromosome Pearson’s r versus the MSE for different loss functions (shown in a different color) and different target resolutions (shown in a different marker). The results for the scaled Pearson’s r loss function was removed due to poor training runs. (**b**) *Bin resolution analysis.* Plot of the whole-chromosome Pearson’s r for models trained on a given bin size (*y*-axis) with predictions that were systematically down-sampled to a lower resolution for evaluation (*x*-axis). (**a**,**b**) Pearson’s r represents the average across cell lines.

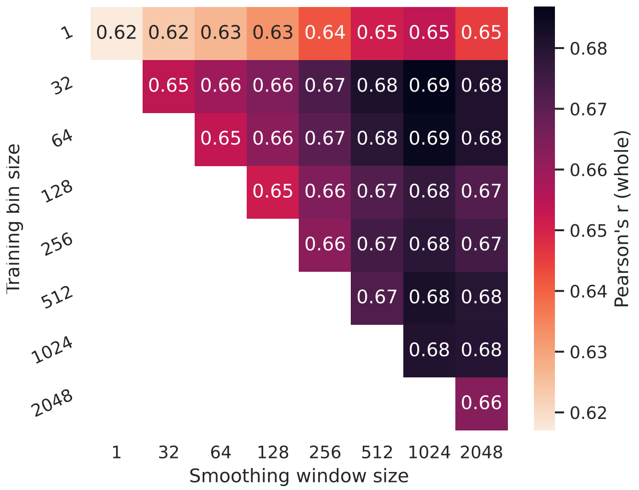


**Extended Data Figure 2.** The effect of smoothing coverage on performance. Basenji-based models were trained on target resolutions (*y*-axis) and evaluated using different levels of smoothing with a box-car filter. For each higher resolution model, a box-car filter was applied to both predictions and experimental coverage values with various kernel sizes prior to calculating the average Pearson's r (*x*-axis). Pearson's r represents the average across cell lines.

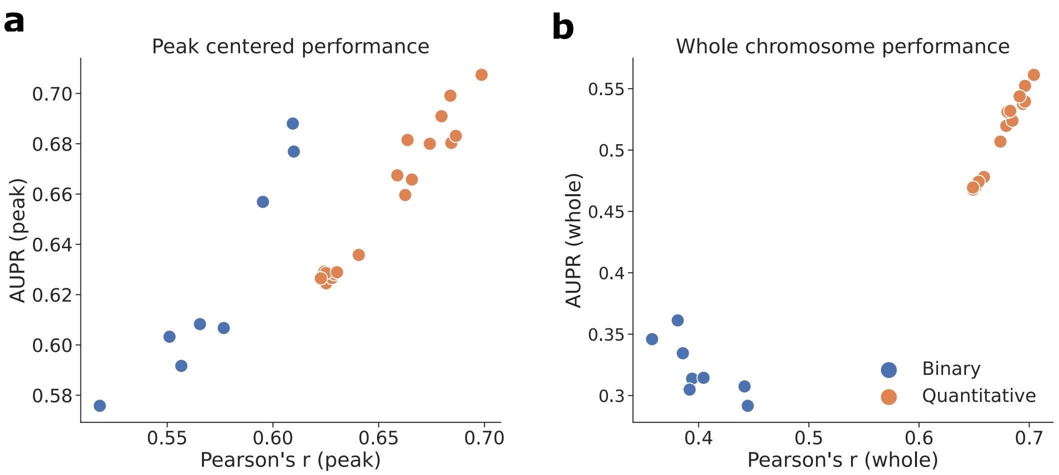


**Extended Data Figure 3.** Performance comparison between quantitative and binary models. Scatter plot of the classification-based AUPR versus the regression-based Pearson's r for various binary models (blue) and quantitative models (orange) on peak-centered test data (left) and whole-chromosome test data (right). Metrics represent the average across cell lines.

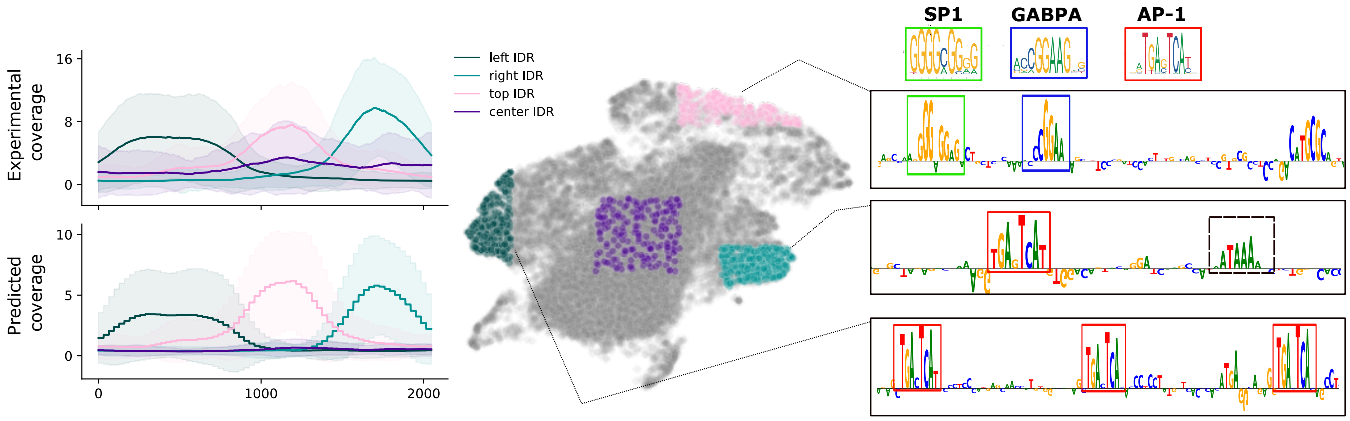


**Extended Data Figure 4.** Interpretability analysis for ResidualBind-32 on PC-3 cell line. UMAP embedding of the penultimate layer representations across all test sequences. (Left) Shows the average experimental coverage and predicted coverage from sequences that map to specific locations in the embedding (shown in a different color). (Right) Representative saliency maps (zoomed in) for sequences within different embedding regions. The known motifs from the JASPAR database are shown at the top and an unknown 'ATAAA' motif is annotated with a box with black dashed lines.

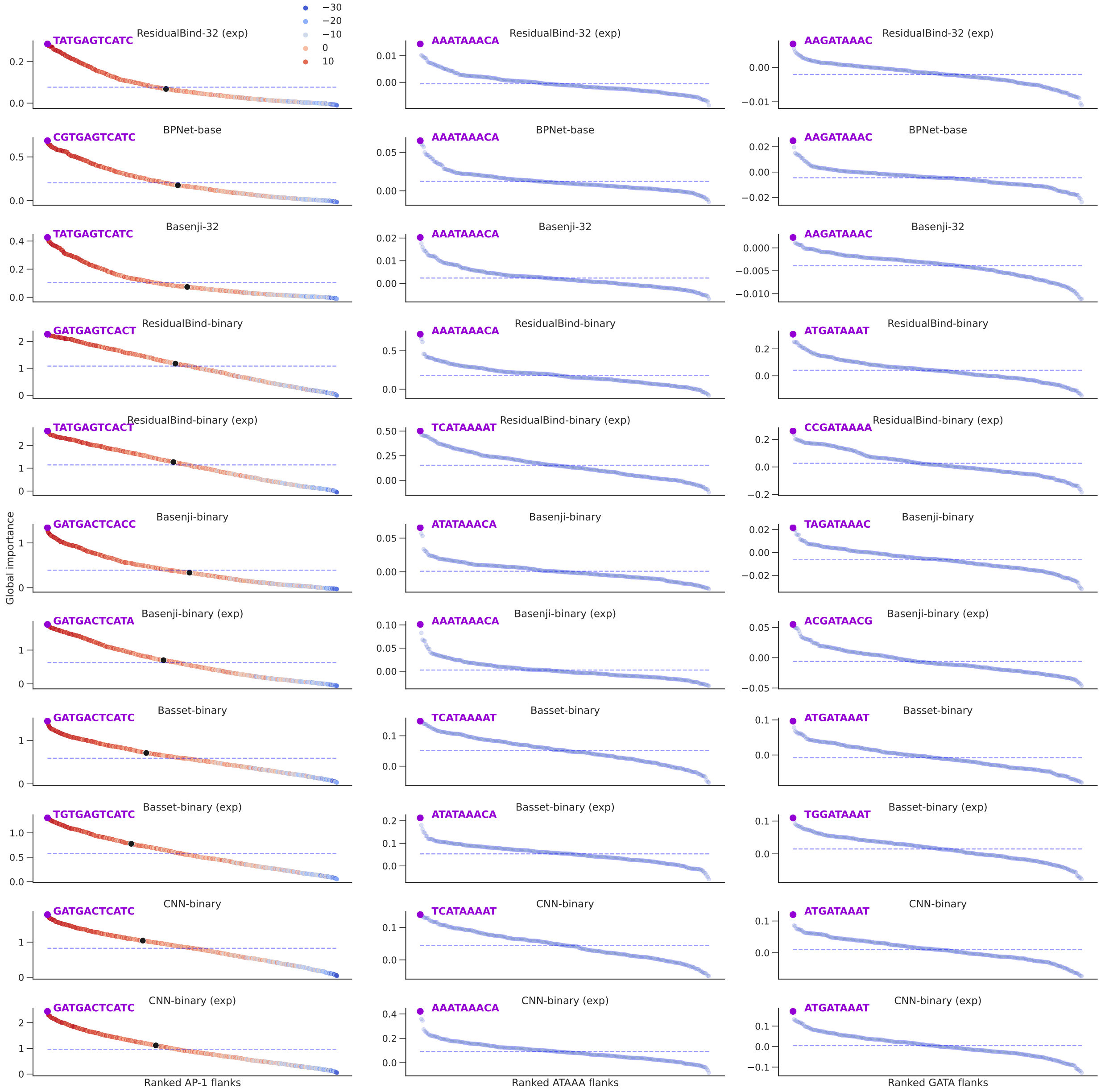


**Extended Data Figure 5.** GIA for optimal flanking nucleotides of motifs in PC-3 cell line for various models. Ranked plot of the global importance for each tested flank for AP-1 motif (left column), ATAAA motif (middle column) and GATA (right column) for different models (shown in a different row). Dashed line represents the global importance of the core motif with random flanks. The hue in the first column represents the position-weight-matrix score for an AP-1 motif from the JASPAR database (ID: MA0491.1). The first 3 rows are quantitative models, the rest are binary models (with (exp) in the name indicating that the first layer ReLU activation has been replaced with an exponential function). For binary models, the results are based on the logits before the output sigmoid activation. The hue in the first column plots represents the PWM score for an AP-1 motif from the JASPAR database (ID: MA0491.1). The black dot in each plot (in the first column) indicates "TGTGATTCATG", which has a high PWM score (12.800) but yields a global importance close to the core motif with randomized flanks.

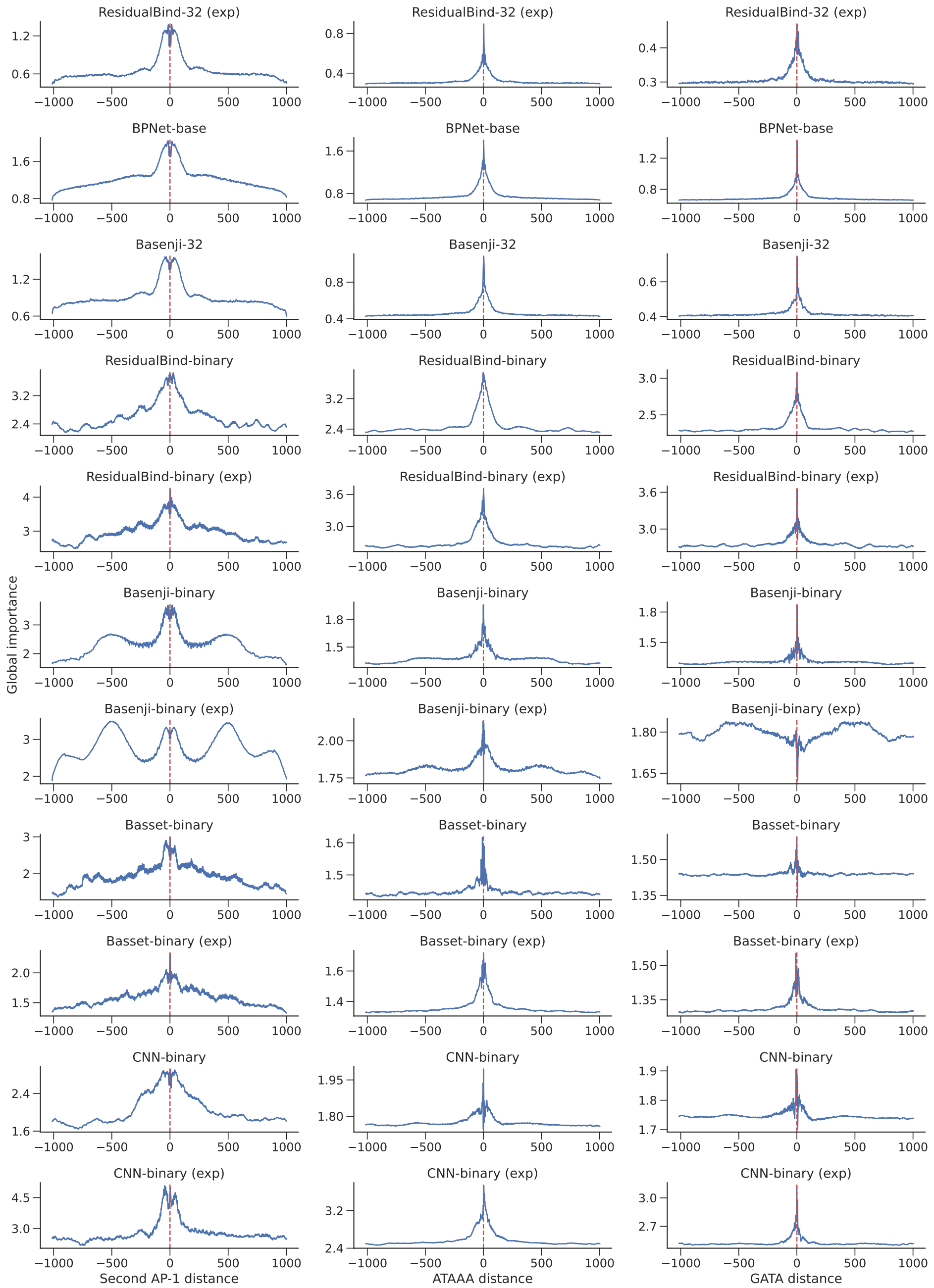


**Extended Data Figure 6.** GIA for distance dependence between AP-1 and other motifs for PC-3 cell line for various models. Global importance plot for sequences with an AP-1 motif fixed at the center of the sequence and another motif that is systematically placed in different locations. Positive and negative values represent the first positions the motifs w/ optimized flanks were embedded to be non-overlapping. First column shows results where the second motif is an identical AP-1 motif, the center column shows results for ATAAA motif and right column for the GATA motif. All the motifs were embedded with optimized flanks. Red vertical dashed lines indicate the 1024bp position. Each row corresponds to a different trained model, the first 3 are quantitative models, the rest are binary models (with (exp) in the name indicating that the first layer ReLU activation has been replaced with an exponential function). For binary models, the results are based on the logits before the output sigmoid activation.

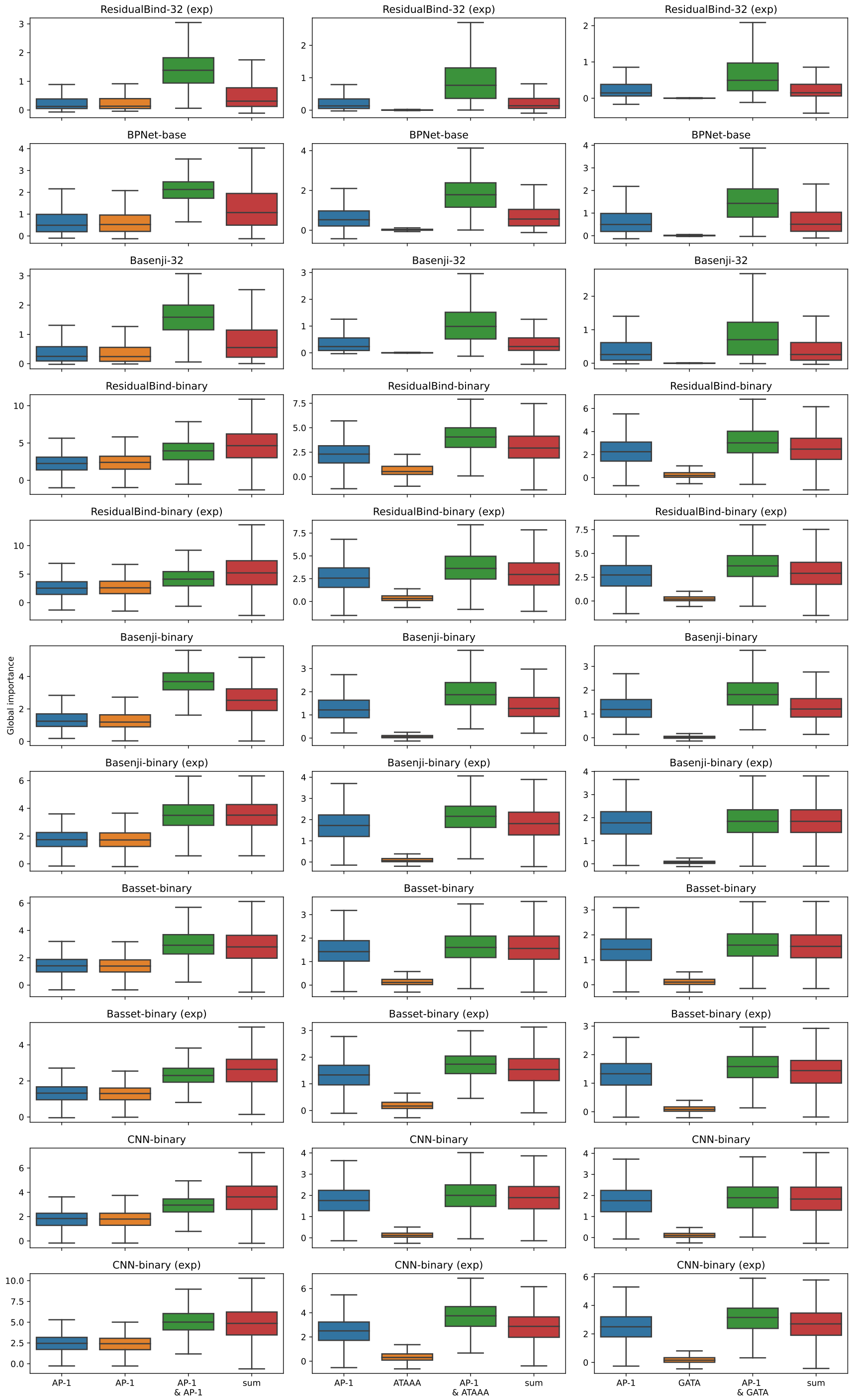


**Extended Data Figure 7.** GIA for cooperative interactions between AP-1 and other motifs for PC-3 cell line for various models. Each column corresponds to a motif pair between two copies of AP-1, ATAAA and AP-1 and AP-1 and GATA. Each row corresponds to a different trained model, the first 3 are quantitative models, the rest are binary models (with (exp) in the name indicating that the first layer ReLU activation has been replaced with an exponential function). For binary models, the results are based on the logits before the output sigmoid activation. Blue and orange box-plots show the global importance scores for the 1000 sampled sequences when motif 1 or motif 2 is individually embedded. Green box-plot shows the case when both motifs are embedded in the same sequence. Red box-plot shows the sum of the green and blue boxes as an estimate of the global importance if there is no interaction. The pairs were embedded at the optimal distance specified from the distance dependence GIA experiments. Box plots show the first and third quartiles, central line is the median, and the whiskers show the range of data with outliers removed.
