## Supplement for "Evaluating deep learning for predicting epigenomic profiles"

**Supplementary Table 1.** Model design and training strategy differences between Basenji and BPNet.

| Features | Basenji | BPNet |
|---|---|---|
| Epigenetic assays | Histone modifications,<br>TF binding,<br>DNA accessibility, 5' cap levels | Binding of 4 TFs |
| Input sequence | 131Kb | 1Kb |
| Target resolution | 128 bin resolution | Base resolution |
| Training set selection | Random selection | Center on IDR peaks |
| Loss function | Poisson NLL | Multinomial NLL + MSE |
| Evaluation Metrics | Pearson's r | Jenson-Shannon divergence |
| Model architecture features | Residual connections,<br>dilated convolutions,<br>maxpool layers | Residual connections,<br>dilated convolutions,<br>task-specific heads |
| Augmentation | reverse complement, random shift (up to 3 bps) | reverse complement |
| Test set selection | Random held out 5% of all sequence data | IDR peaks on held-out test chromosomes 1, 8 and 9 |

**Supplementary Table 2.** CAGI5 replicability. Each CAGI saturated mutagenesis experiment was conducted for three replicates. Pearson correlation across replication results were measured to estimate experiment consistency.

| Name | Replicate correlation |
|---|---|
| F9 | 0.61 |
| FP1BB | 0.74 |
| HBB | 0.62 |
| HBG1 | 0.78 |
| HNF41 | 0.75 |
| IRF4 | 0.98 |
| IRF6 | 0.90 |
| LDLD | 0.99 |
| MSMB | 0.75 |
| MYC | 0.55 |
| PKLR | 0.79 |
| SORT1 | 0.98 |
| TERT(GBM) | 0.90 |
| TERT(HEK293T) | 0.65 |
| ZFAND3 | 0.72 |

**Supplementary Table 3.** ATAC-seq experiments selected from ENCODE database. The IDR-peak bed file and log-fold over control BigWig files were taken based on the following ENCODE experimental accessions.

| Cell line | Experiment accession |
|---|---|
| GM21381 | ENCSR512YXO |
| GM23338 | ENCSR485TLP |
| HepG2 | ENCSR291GJU |
| RWPE2 | ENCSR080SNF |
| HG03575 | ENCSR331JFZ |
| K562 | ENCSR868FGK |
| DND-41 | ENCSR660WSB |
| GM12878 | ENCSR637XSC |
| A549 | ENCSR032RGS |
| HCT116 | ENCSR872WGW |
| IMR-90 | ENCSR200OML |
| NCI-H929 | ENCSR382LBS |
| Panc1 | ENCSR591PIX |
| PC-3 | ENCSR499ASS |
| MCF-7 | ENCSR422SUG |

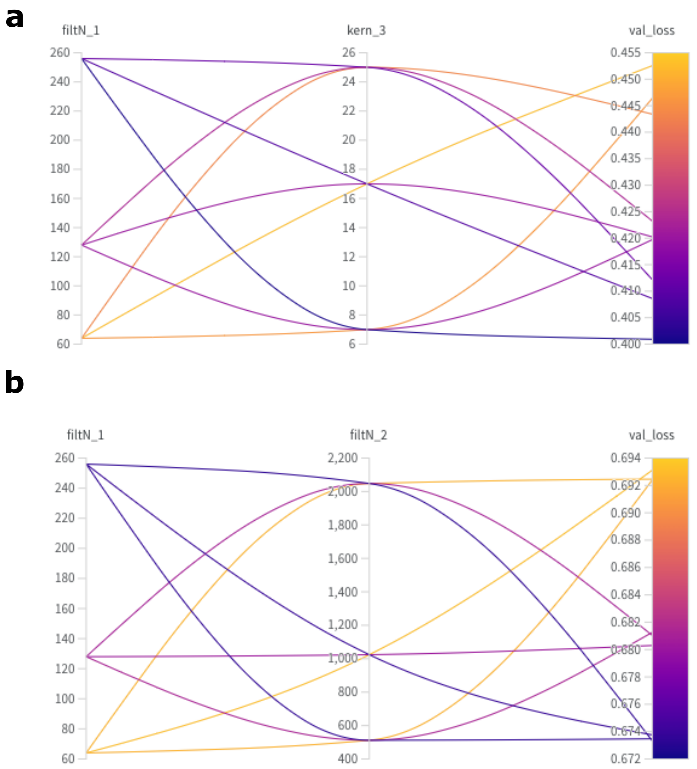


**Supplementary Figure 1.** Hyperparamter search with WandB. Parallel plot for different hyperparameters used in a grid search for (**a**) BPNet-base and (**b**) Basenji. (**a**) BPNet-base optimized hyperparameters were the number of kernels in the first convolutional layer (filtN_1), which is then fixed throughout all subsequent convolutional layers, and the kernel size (kern_3) of the task-specific output heads. (**b**) Basenji-based optimized hyperparameters specifying the number of filters in the first convolutional layer (filtN_1) and the second convolutional layer (filtN_2), which correspondingly sets the number of filters in the subsequent residual blocks.

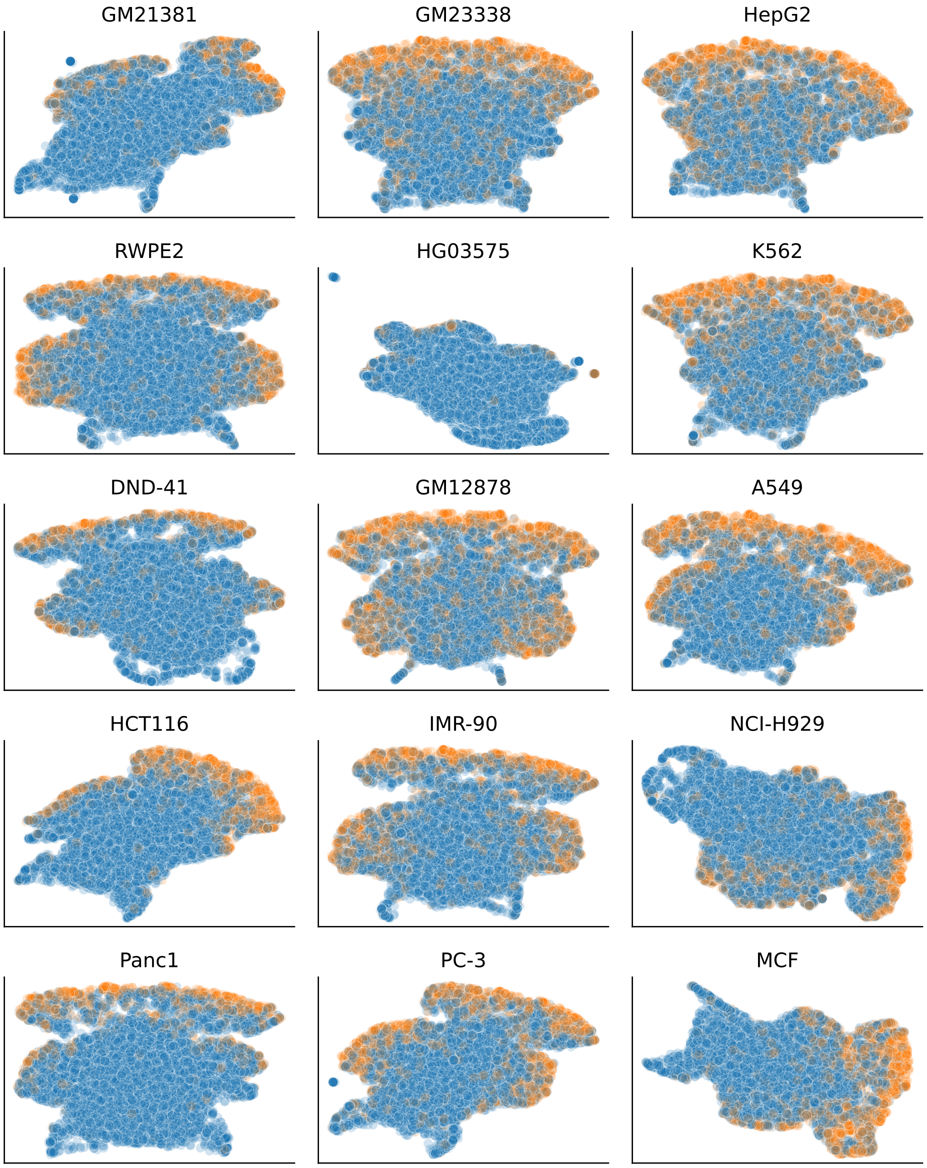


**Supplementary Figure 2.** UMAP embeddings of the test set representations for Residualbind-32 with single-task outputs and exponential activations. For each cell line, embedded sequences were selected based on the coverage value above a threshold of 2. Orange dots indicate sequences that overlap with a statistically significant peak.

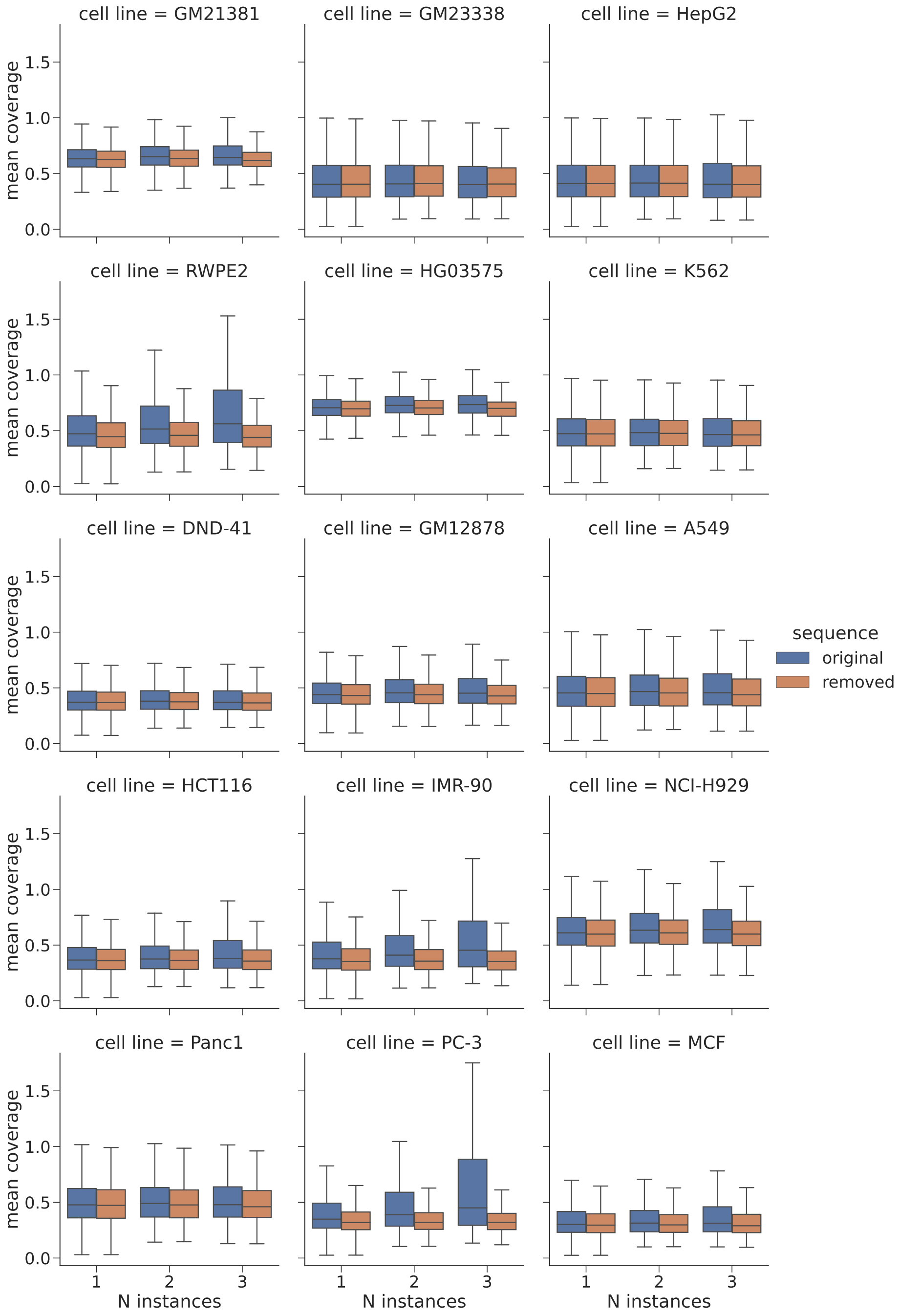


**Supplementary Figure 3.** Occlusion analysis for AP-1 motifs using Residualbind-32 with single-task outputs and exponential activations. Box-plots for the mean coverage value of 10,000 randomly sampled sequences from the test set that contained 1, 2, and 3 (or more) AP-1 motifs and the mean coverage values when the AP-1 motifs are replaced by randomized sequences (averaged across 20 random trials) for each cell line. 3,381, 1,310, and 323 sequences contained 1, 2, and 3 instances of the AP-1 motif, respectively. Box plots show the first and third quartiles, central line is the median, and the whiskers show the range of data with outliers removed.
